# Comparative transcriptional analysis of the satellite glial cell injury response

**DOI:** 10.1101/2021.11.22.469443

**Authors:** Sara Elgaard Jager, Lone Tjener Pallesen, Lin Lin, Francesca Izzi, Alana Miranda Pinheiro, Sara Villa-Hernandez, Paolo Cesare, Christian Bjerggaard Vaegter, Franziska Denk

**Affiliations:** Wolfson Centre for Age-Related Diseases, Institute of Psychiatry, Psychology and Neuroscience, King’s College London, Guy’s Campus, London, United Kingdom; Department of Biomedicine, Danish Research Institute of Translational Neuroscience—DANDRITE, Nordic-EMBL Partnership for Molecular Medicine, Aarhus University, Aarhus C, Denmark; Department of Biomedicine, Aarhus University, Denmark & Steno Diabetes Center Aarhus, Denmark; NMI Natural and Medical Sciences Institute at the University of Tübingen, Tübingen, Germany

## Abstract

Satellite glial cells (SGCs) tightly surround and support primary sensory neurons in the peripheral nervous system and are increasingly recognized for their involvement in the development of neuropathic pain following nerve injury. The SGCs are difficult to investigate due to their flattened shape and tight physical connection to neurons *in vivo* and their rapid changes in phenotype and protein expression when cultured *in vitro*. Consequently, several aspects of SGC function under normal conditions as well as after a nerve injury remain to be explored. The recent advance in single cell RNAseq technologies has enabled a new approach to investigate SGCs. Here we publish a dataset from mice subjected to sciatic nerve injury as well as a dataset from dorsal root ganglia cells after 3 days in culture. We use a meta-analysis approach to compare the injury response with that in other published datasets and conclude that SGCs share a common signature following sciatic nerve crush and sciatic ligation, involving transcriptional regulation of cholesterol biosynthesis. We also observed a considerable transcriptional change when culturing SGCs, suggesting that some differentiate into a specialised *in vitro* state, while others start resembling Schwann cell-like precursors. The datasets are available via the Broad Institute Single Cell Portal.

## Introduction

Satellite glial cells (SGCs) are located in peripheral ganglia, where they tightly envelop each neuronal cell body into defined SGC-neuron units (Ennio Pannese, 1981, 2010). With their flattened morphology and only approximately 20μm distance from the neuronal soma, they are ideally located to communicate with neurons and provide a protected homeostatic microenvironment. Accordingly, in the healthy organism, SGCs have been shown to maintain the extracellular space by buffering glutamate and K^+^ concentrations (Duce & Keen, 1983; Miller, Richards, & Kriebel, 2002; J. P. Vit, Jasmin, Bhargava, & Ohara, 2006; Procacci, Magnaghi, & Pannese, 2008), and they have furthermore been shown to modulate neuronal excitability via bi-directional ATP signalling (Chen et al., 2008; Lemes et al., 2018) and AMPA and NMDA receptors (Kung et al., 2013).

Several studies have investigated the responsiveness of SGCs in rodent models of nerve injury, where the peripheral axonal branch is damaged through e.g. ligation, transection or crush. Despite such neuronal injury being induced at a substantial distance from SGCs in the dorsal root ganglia (DRG), it clearly has a knock-on effect on their function (Avraham et al., 2020; Jager et al., 2020; Menachem Hanani & Spray, 2020; Avraham, Feng, et al., 2021). To date, SGC reactivity has mainly been studied with focus on changes in ATP signalling between neurons and SGCs, a decrease in K^+^ buffering capacity, and an increase in the number of SGC-SGC gap junctions. Thus, somata of injured neurons are believed to release ATP in an action potential dependent manner, activating P2Y4, P2X7 and/or P2Y12 receptors on SGCs. This, in turn, modulates feedback signalling and, ultimately, the excitability of neurons (Weick et al., 2003; J. P. Vit et al., 2006; Zhang, Chen, Wang, & Huang, 2007). A decreased K^+^ buffering of SGCs is thought to be driven by a reduced expression of the Kir4.1 channel (J.-P. Vit, Ohara, Bhargava, Kelley, & Jasmin, 2008; Tang, Schmidt, Perez-Leighton, & Kofuji, 2010; Takeda, Takahashi, Nasu, & Matsumoto, 2011). This likely contributes to an increased concentration of extracellular K^+^ within the SGC-neuron unit and thereby increases neuronal excitability (J. P. Vit et al., 2006). Finally, changes are observed in SGC-SGC gap junction connectivity, with a rise in the expression and functional assembly of connexin43 (M. Hanani, Huang, Cherkas, Ledda, & Pannese, 2002; E Pannese, Ledda, Cherkas, Huang, & Hanani, 2003; Cherkas et al., 2004; Huang, Cherkas, Rosenthal, & Hanani, 2005; Dublin & Hanani, 2007; Suadicani et al., 2010). While such increased gap junction connectivity has been shown to be important for facilitating the spread of Ca^2+^ waves (Suadicani et al., 2010), the functional consequences of this in relation to SGC-neuron communication remains unclear.

Relatively little is still known about the basic biology of SGCs, primarily due to their flattened morphology and close proximity to neurons which complicates immunohistochemical studies and *in vivo* experiments (Ennio Pannese, 2010). Additionally, SGCs rapidly change their phenotype in culture, making *in vitro* experiments similarly challenging (Belzer, Shraer, & Hanani, 2010; George, Ahrens, & Lambert, 2018). It is therefore encouraging that recent advances in single cell RNA sequencing (scRNAseq) have made it possible to study the transcriptional profile of these cells in previously unprecedented detail. To date, six papers and manuscript preprints have included such SGC scRNAseq studies in mice with focus on either development (Mapps et al., 2021; Tasdemir-Yilmaz et al., 2021), species comparison (Avraham et al., 2021) or nerve injury (Avraham et al., 2020; Renthal et al., 2020; K. Wang et al., 2021). Furthermore, our team published a bulk RNA-seq experiment with focus on nerve injury (Jager et al., 2020).

Here, we present two additional scRNAseq datasets on mouse SGCs. We analysed our novel datasets in conjunction with those previously published, to investigate whether transcriptional injury responses are common and reproducible across models and laboratories. While we were able to identify a reproducible transcriptional nerve injury signature in SGCs, the number of genes found commonly regulated across datasets was small. Furthermore, we compare the transcriptional profiles of mouse DRG SGCs acutely isolated with those cultured *in vitro*. Our findings confirm that cultured SGCs indeed present a very different transcriptional profile relative to those acutely isolated (George et al., 2018). The datasets and analyses are compiled and accessible at the Broad Institute’s Single Cell Portal (www.singlecell.broadinstitute.org/single_cell/study/SCP1539/) for further investigations of genes of interest.

## Methods

### Animals

All mice were housed under standard conditions with 12h light/dark cycle and free access to standard chow and water. For the spared nerve injury (SNI) experiment, 13-week-old male C57BL/6J mice from Janvier labs were housed in pairs of 2 littermates. Small-grained bedding was used after SNI. The SNI experiment was approved by the Danish Animal Experiments Inspectorate under the Ministry of Environment and food (permission number 2017-15-0201-01192-C1). For the culture experiment, 2-week-old male and female SWISS mice from Janvier labs were used. This animal experiment was conducted in accordance with the EU legislation for the care and use of laboratory animals (Directive 2010/63/EU) and the German Animal Welfare Act (“Tierschutzgesetz”, 2019).

### Spared nerve injury (SNI)

SNI was performed in the left and right hindleg according to the method described previously (Richner, Bjerrum, Nykjaer, & Vaegter, 2011). The procedure was performed under isoflurane (IsoFlo Vet, Abbott) anaesthesia. The sciatic nerve was exposed with skin incision and blunt dissection of the overlying muscle. A 6.0 vicryl suture was used to tightly ligate and then cut the common peroneal and tibial branches of the sciatic nerve, with the sural nerve left intact. The wound was closed with surgical tissue adhesive (Indermil Tissue Adhesive, Henkel), and for local analgesia a droplet of lidocaine SAD (10 mg/ml; Amgros I/S) was applied to the wound. Buprenorphine (0.3 mg/ml; Temgesic, RB Pharmaceuticals) and the antibiotic ampicillin (250 mg/ml; Pentrexyl; Bristol-Myers Squibb) were mixed and diluted 1:10 in isotonic saline (9 mg/ml; Fresenius Kabi) and 0.1 ml was injected subcutaneously following surgery for peri-operative analgesia and protection against infection. The operation was performed bi-laterally to ensure enough material for the sequencing and eight L3 and L4 DRGs were collected from 2 mice per condition (naïve, 7 days post SNI and 14 days post SNI).

### Cultured DRG cells

The 2-week-old mice were euthanized with CO_2_ before they were disinfected in 70% ethanol and decapitated. DRGs from cervical, thoracic, and lumbar levels were dissected. The ganglia were then incubated in 2.5 ml CD dissociation buffer (DMEM + GlutaMAX, Thermo Fisher, 31966-021 with 3.6 mg/ml glucose, Carl Roth, NH06.3, 3 mg/ml Collagenase type IV, Worthington, LS004186, and 6 mg/ml Dispase, Worthington, LS02109) for 40 min at 37°C. Next, the CD dissociation buffer was replaced by 5 ml D dissociation buffer (DMEM + GlutaMAX with 3.6 mg/ml glucose, and 3mg/ml Dispase) for a further 40 min at 37°C. Following enzymatic digestion, the cells were manually triturated in cell medium (DMEM + GlutaMAX with 5% horse serum, Thermo fisher, 26050-070 and 0.5% Penicillin-Streptomycin, Sigma Aldrich, P4333-20ML), after which the cellular solution was cleared of debris by gradient centrifugation through 4 layers of various percentages of OptiPrep™ (Sigma Aldrich, D1556-250ML) in cell medium (from the bottom: 28% OptiPrep with resuspended cells, 15%, 8% and 0%). The gradient was centrifuged at 800xg for 22 min, and cells were recovered from the interface between the 15% and 8% layer. The cells were plated on laminin coated 24-well plates and kept for 72h at 37°C, 5% CO_2_.

### Dissection and processing of DRGs for scRNAseq

#### SNI experiment (Cell_SNI)

Mice were anaesthetized using isoflurane and transcardially perfused using 10-20 ml DPBS (Thermo Scientific, SH3002802). L3 and L4 DRGs were identified and collected from both sides as previously described (Richner, Jager, Siupka, & Vaegter, 2017). For each time point (naïve, 7 days and 14 days post SNI) L3 and L4 DRGs were dissected from 2 mice (8 DRGs/sample) and stored in ice-cold HBSS (Gibco, 14170088). DRGs were centrifuged for 4 min at 500 xg at 4°C and incubated in 1 ml dissociation buffer (2.5mg/ml collagenase, Sigma Aldrich, C9722, and 5 U/ml dispase II, Sigma Aldrich, D4693 in DPBS) for 30 min at 37°C in 5% CO_2_. Following enzymatic digestion, the cells were manually triturated using a p1000 pipette until homogenous. 9 ml of PBS was added (Sigma, D8537), and cells were centrifuged at 500xg for 8 min, 4°C and incubated for 10 min at 37°C in 0.5ml trypsin-EDTA (0.25% trypsin w/v and 0.1% EDTA w/v, Sigma 59418C diluted 1:1 in DPBS). 5 ml HBSS with 10% (v/v) Fetal Bovine Serum (FBS, Sigma, F9665) was added to stop the reaction. The cell suspension was centrifuged at 500xg, 4°C for 8 min, and the cell pellet resuspended in 1 ml HBSS with 40 Kunitz units Deoxyribonuclease I (Sigma Aldrich, DN25-1G) before filtration through a 40µm cell strainer (VWR, 734-0002). Following a final centrifugation for 10 min at 500 xg, 4°C the cells were resuspend in PBS, 5% (v/v) FBS at a concentration of 1000 cells/ul.

#### Cultured DRG cells (Cell_culture)

The cells were maintained in culture for 72h before they were detached with Trypsin-EDTA 0.25% w/v, centrifuged, counted, and processed for 10X scRNAseq.

### scRNAseq on 10X Chromium (Cell_SNI and Cell_Culture)

To construct scRNAseq libraries, the cell suspensions were processed with the Chromium Single Cell 3’ GEM, Library & Gel Bead Kit v3 (10x Genomics, PN 1000075) according to the manufacturer’s instructions. During this process, the 10X Chromium device uses a microfluidic system to partition each cell into a single droplet, each containing sequencing barcodes and the enzymes required for reverse transcription. The barcodes are specific to each droplet and ensure that it is possible to identify which transcripts were detected in which cell after sequencing. The libraries were sequenced using DNBSEQ-G400. The raw sequencing reads were processed using Cell Ranger version 3.0.2 and mapped to the reference genome mm10-3.0.0, Ensemble 93. The three Cell_SNI conditions (naïve, 7 days, 14 days) were processed on the same 10X chip and in the same subsequent library preparation, in order to minimise batch effects.

### Quality control, clustering and visualization of scRNAseq

The Cell_SNI and Cell_culture count matrices were analysed with Seurat v3 (Stuart et al., 2019) in R. The previously published datasets with focus on nerve injuries (Avraham et al., 2020; Renthal et al., 2020; K. Wang et al., 2021) were reanalysed with Seurat v3 from the count matrices made available on the Gene Expression Omnibus website (GSE139103, GSE154659 and GSE155622). To ensure that we analyse high quality cells, we started by filtering out those with less than 200 detected genes. We further filtered out likely dead cells, based on mitochondrial gene expression permitting a maximum of 30% gene expression being mitochondrial, tailoring the precise cut-off value to each individual dataset (Table 1).

**Table 1:** Overview of details for analysis of the datasets.

| Dataset | Cut-off for mitochondrial genes (%) | Resolution for clustering | Dimensions for PCA and clustering | Doublet cut-off for SGC cluster (# nFeature_RNA) |
| --- | --- | --- | --- | --- |
| Cell_SNI | 30% | 0.08 | 1:20 | 3500 |
| Cell_crush | 15% | 0.08 | 1:20 | 2500 |
| Cell_culture | 10% | 0.18 | 1:20 | NA |
| Renthal et al. | 10% | 0.5 | 1:20 | NA |
| Wang et al. | 5% | 0.08 | 1:20 | NA |

Next, we performed normalization of the raw transcript counts detected in each cell. The normalization is a two-step process consisting of scaling and transformation. Scaling is performed by calculating counts per 10,000 counts. This provides count concentration instead of absolute number, which is useful since cells vary in size and therefore also in number of mRNA molecules. Furthermore, scaling removes efficiency noise, which arises because the v3 chemistry used for sequencing is not equally effective in each droplet. Next, natural-log transformation using log1p is performed on each scaled count number for each gene in each cell. This ensures that highly expressed genes are not given more weight in the downstream integration analysis compared to lowly expressed genes. Following normalization, we identified the 2000 genes that showed the highest cell-to-cell variability. Where necessary (i.e. when comparing across 10X chips/ experiments from different laboratories) we used these genes to integrate across conditions. Integration mitigates the impact of batch effects on subsequent cluster analysis (Tran et al., 2020). We next applied a linear transformation step and performed PCA, which is used as the foundation for clustering and Uniform Manifold Approximation and Projection (UMAP, Table 1). UMAP provides a two-dimensional reduction, enabling visualization of the datasets, while the clustering identifies similar cells. Finally, we identified the genes that are highly expressed in each cluster (marker genes) and used those to annotate the clusters with a cell type (Supplementary notebook).

### Comparison of dataset annotations with scMAP

The annotations of the clusters in the Cell_crush dataset from Avraham *et al*. (Avraham et al., 2020) and the Cell_SNI dataset were compared to each other with scMAP (Kiselev, Yiu, & Hemberg, 2018) to ensure annotation consistency. We used scMAP to project each cell in the Cell_SNI dataset to the cell types identified in the Cell_crush dataset. The projection is based on the 500 most informative genes identified with the selectFeatures function in the scMAP package (Supplementary notebook). The selectFeatures function use a linear model to capture the relationship between mean expression and number of dropouts (zero expression). The most informative genes are identified as the ones with more dropouts than expected, i.e. those not present in some clusters. The output of the scMAP projection is a Sankey plot illustrating how the annotations in the datasets compare to each other.

### Additional quality control of the SGC cluster

With droplet-based sequencing technologies like 10X, there is a risk of duplets, with two cells being captured in the same droplet, and barcoded as one. An often-used strategy to eliminate duplets, is to set a threshold for the number of detected genes in each cell. However, the cell types contained within a DRG are very heterogenous, ranging from very large sensory neurons to small immune cells. The difference in cell size results in the detection of relatively many genes in neurons and fewer genes in the immune cells (see Supplementary Figure 1) (Padovan-Merhar et al., 2015). Due to this heterogeneity, it is not possible to set a universal threshold that filters out SGC duplets without depleting neurons. For all our downstream analyses focused on SGCs only, we therefore subset the SGC cluster and adjusted our duplet-filtration threshold to fit this particular cell population (Supplementary notebook and Table 1).

### Differentially expressed genes in SGCs

Differentially expressed genes were identified with Seurat v3 based on the unintegrated data using the non-parametric Wilcoxon rank sum test. For us to consider a gene to be differentially expressed, it needed to be expressed in at least 10% of SGCs in either the naïve or injured conditions, have a log2 fold change of at least 0.25 and an adjusted p value of less than 0.05. To avoid introducing technical artefacts, we only performed these analyses within individual batch-controlled datasets (Cell_SNI and Cell_crush; i.e. those deriving from the same 10X chip) and then compared the resulting lists of differentially expressed genes across studies. The gene ontology enrichment was done with Metascape (Zhou et al., 2019) using their web interface for multiple lists.

### Comparison of isolated and cultured SGCs (Cell_SNI versus Cell_culture)

The Cell_SNI and Cell_culture datasets including all conditions were integrated using Seurat v3. Joint clusters were identified and annotated as described above. To investigate the SGC cluster further, we subset it and performed normalization, integration, clustering and visualization again on the raw counts. This resulted in 5 different SGC subclusters. We compared the transcriptome of our joint SGC dataset to a scRNAseq dataset of the developing mouse nervous system from Furlan *et al*. (Furlan et al., 2017), using the matchReferences() function of SingleR (Aran et al., 2019) (see Supplementary notebook for more details). The function finds the probability of a cell in the SGC dataset being assigned each label in the dataset from Furlan *et al*. and vice versa. A probability of 1 indicates that there is a 1:1 relation between that pair of labels while a probability of 0 indicates that the cell clusters are not similar.

## Results

### Cell annotations in different scRNAseq data sets

In this analysis four different sets of single cell or single nucleus RNAseq data from mouse DRGs after different nerve injuries (see Table 2) were included. Three datasets are published (Avraham et al., 2020; Renthal et al., 2020; K. Wang et al., 2021) and available online (GSE139103, GSE154659 and GSE155622) while the fourth scRNAseq dataset of SNI responses at day 7 and 14 in the DRG is published in this work (GSE174430). The overall goal of this analysis was to identify if SGCs share a common response to nerve injuries across different experimental conditions.

**Table 2:** Overview of the different single cell and single nucleus RNAseq datasets analysed in this paper.

| Publication | Dataset | Mouse strain | Age | Sample prep | Condition | scRNA-seq | Injury analysis | Culture comparison analysis |
| --- | --- | --- | --- | --- | --- | --- | --- | --- |
| Avraham et al. 2020 | Cell_crush | C57Bl/6J | 8-12 weeks | Dissociation with collagenase and papain, sorted with FACS | Crush, 3 days | Whole cell, 10X | Yes | No |
| N/A | Cell_SNI | C57Bl/6J | 13 weeks | Dissociation with collagenase and dispase, followed by trypsin | SNI, 7 & 14 days | Whole cell, 10X | Yes | Yes |
| N/A | Cell_culture | SWISS | 2 weeks | Dissociation with collagenase and dispase | uninjured, cultured for 3 days | Whole cell, 10X | No | Yes |
| Renthal et al. 2020 | N/A | C57Bl/6J | 8-12 weeks | Extraction of nuclei | various injuries | Nucleus, InDrops | No | No |
| Wang et al. 2021 | N/A | C57Bl/6J | 7-8 weeks | Dissociation with enzymes, sorted with Percoll gradient | SNI at various time points | Whole cell, 10X | No | No |

First, the published datasets were re-analysed, focusing specifically on SGC clusters. It was apparent that the SGC scRNAseq data from Renthal *et al*. and Wang *et al*. contain a substantial amount of neuronal background signal. Specifically, the top differentially expressed genes after nerve injury in non-neuronal cell types all resemble the same ‘canonical’ neuronal response profile. For example, genes such as Gal, Atf3, Npy, Nts and Sprr1a were regulated in the SGC clusters (see Supplementary Figure 2 and Excel Sheets “Renthal et al 7d” and “Wang et al 7d” for the full list of differentially expressed genes). While both studies contain an impressive amount of data, with cells taken from many different time points and nerve injury models, both were also designed with a focus on DRG neurons – and as it turns out, this impacts their suitability for the analysis of differential gene expression in non-neuronal cells. To avoid any bias in our SGC analysis, these datasets therefore had to be excluded from the meta-analysis.

From the two remaining datasets, the sciatic nerve crush data from Avraham *et al*. (Cell_crush) (Avraham et al., 2020) and the sciatic nerve ligation dataset (Cell_SNI) presented here, different cell populations were identified using unsupervised clustering, visualised with UMAP plots (Figure 1A). To determine the nature of each cell cluster, the expression of marker genes was investigated (Figure 1B) and the datasets were annotated individually based on the top marker genes for each cell type (Figure 1A). The annotations were shown to be consistent between datasets using the package scMAP, which projects one dataset annotation on to the other (Figure 1C). No differences in cell types present in the datasets were detected, however, minor variations in the cell proportions were observed. Specifically, more Schwann cells were detected in Cell_SNI, and more fibroblasts and macrophages in Cell_crush (Figure 1D). This phenomenon is presumably due to the different dissociation techniques applied (Table 2).

**Figure 1:**
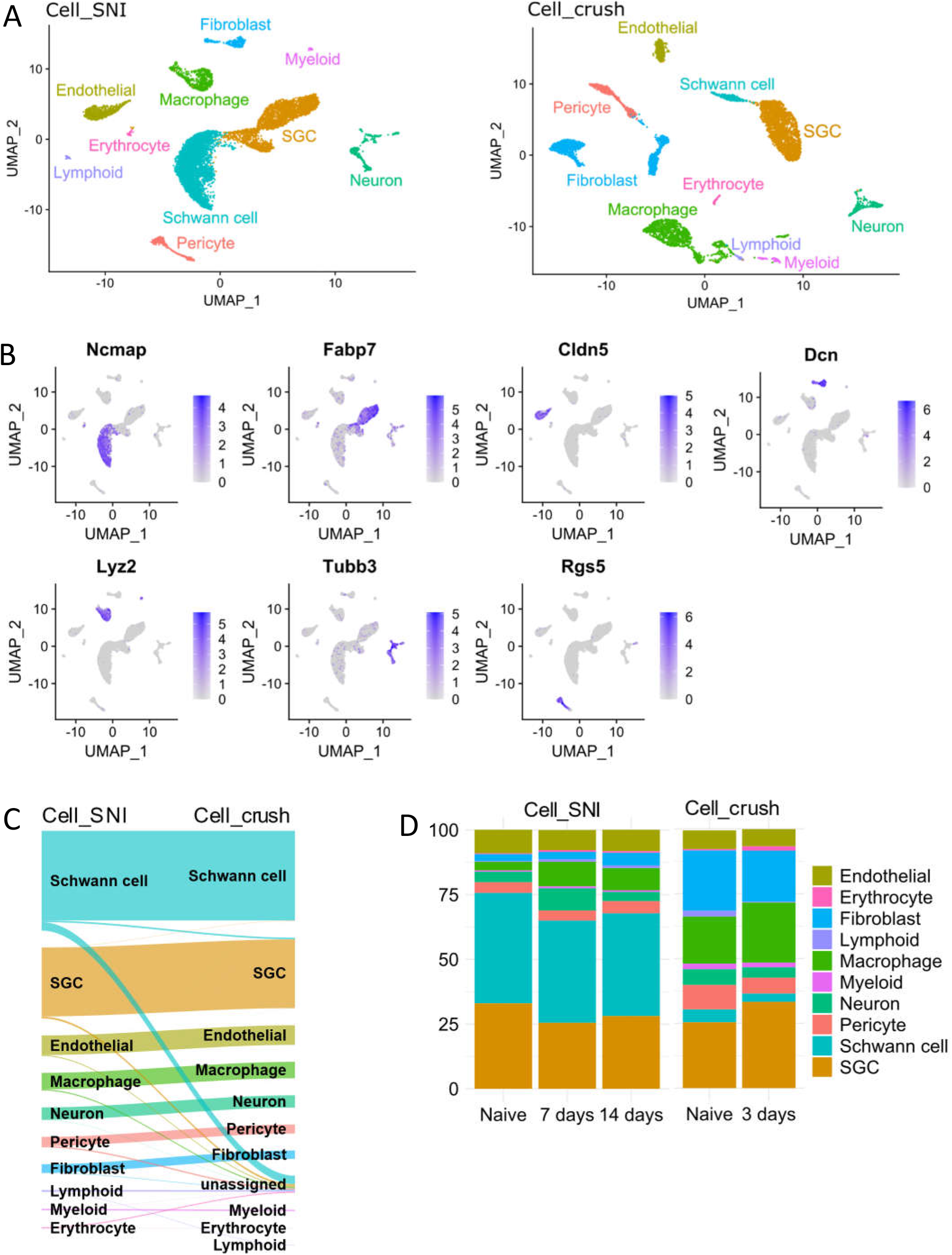
SGCs are easily identifiable in the Cell_SNI and Cell_crush datasets. A) UMAPs of the Cell_SNI and Cell_crush datasets highlighting the identified cell types. B) UMAPs of the Cell_SNI dataset highlighting gene expression used to identify the cell types. Ncmap = Schwann cells, Fabp7 = SGCs, Cldn5 = endothelials, Dcn = fibroblasts, Lyz2 = macrophages, Tubb3 = neurons, Rgs5 = pericytes. C) Sanky diagram showing the projection of the Cell_SNI dataset on to the Cell_crush dataset. D) The percentage distribution of the cell types in the dataset. Cell_SNI naïve = 3486 cells, Cell_SNI 7 days = 3029 cells, Cell_SNI 14 days = 4386 cells, Cell_crush naïve = 3090 cells, Cell_crush 3 days = 3748 cells

### SGCs demonstrate a common response to nerve injury across tested conditions and timepoints

The response of SGCs to nerve injury was investigated. Both datasets contain SGCs from L3-L5 DRGs following sciatic nerve injury. Many, but not all, neuronal somata in these DRGs project their axons to the sciatic nerve (Rigaud et al., 2008). Consequently, not all SGCs in the samples from injured conditions would have been surrounding an injured neuron.

First, it was assessed if unsupervised cluster analysis could distinguish SGCs that had been surrounding an injured neuron from those that had not. To ensure that the analysis contained enough data to enable sub-clustering, the SGCs from the two datasets were combined and integrated (Figure 2A). It was, however, not possible to identify a cluster consisting exclusively of SGCs from injured mice, neither when analysing all SGCs integrated together (Figure 2B and C), nor when analysing them individually within each dataset (Supplementary Figure 3). This suggests that the differences induced by nerve injury are comparatively more subtle in SGCs than in DRG neurons, which clearly cluster together when damaged (Renthal et al., 2020).

**Figure 2:**
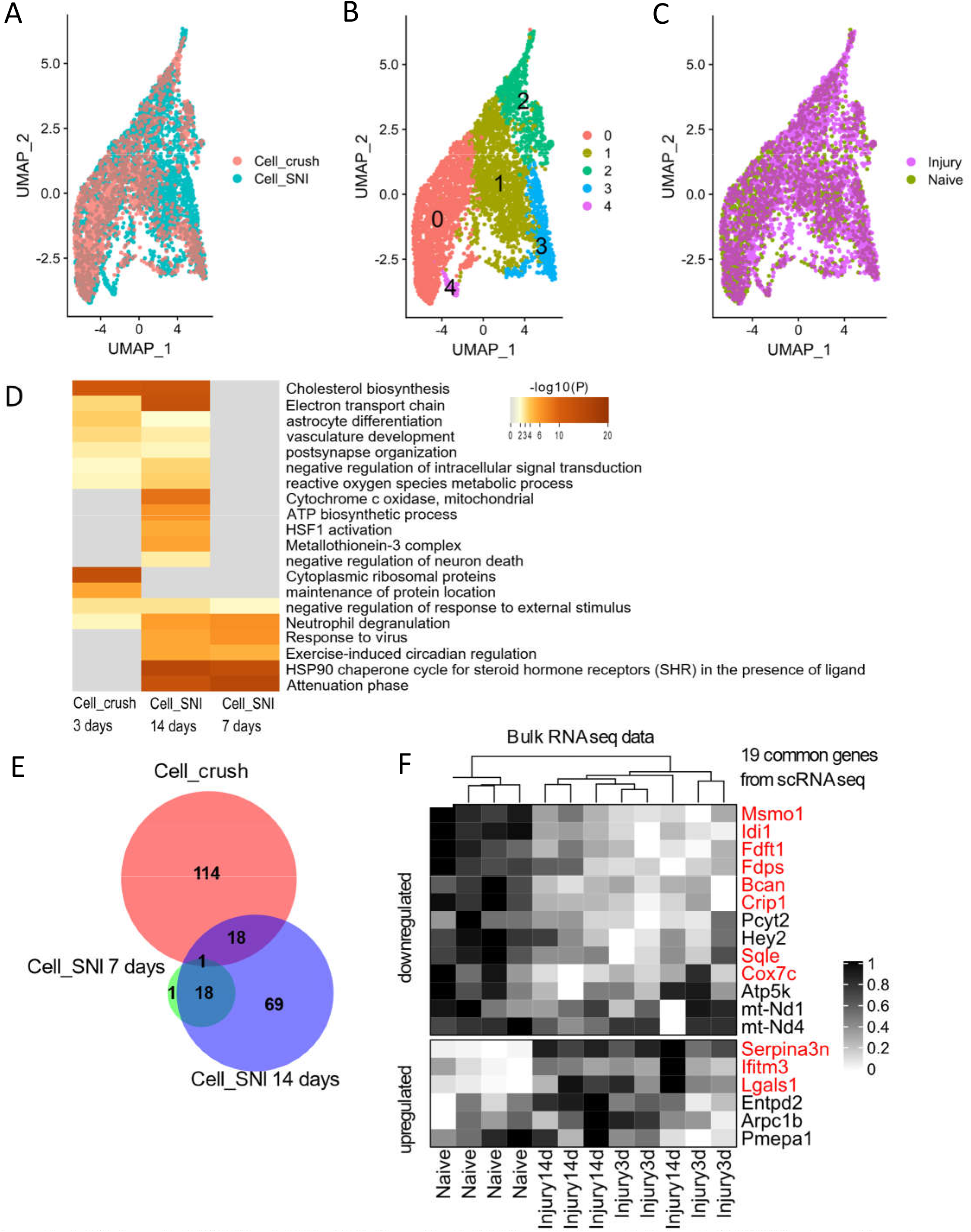
A-C) Integrated UMAPs of all the SGCs from the Cell_SNI and Cell_crush dataset. A) UMAP coloured based on dataset. B) UMAP coloured based on clustering. C) UMAP coloured based on injury condition. No cluster contains only injured SGCs. D) Heatmap of enriched gene annotation terms. The top 20 highest ranking terms are shown. E) Venn diagram of number of differentially expressed genes in SGCs in Cell_crush and Cell_SNI datasets when comparing injured states to naïve. F) Heatmap displaying expression levels from bulk RNAsea data (Jager et al) containing n=4 for per condition (naïve, 3 days and 14 days after injury). The genes extracted here are the 18+1 common genes between the Cell_SNI and Cell_crush datasets from figure 2C. The genes marked in red are also differentially regulated in the displayed bulk RNAseq.

Next, differentially expressed genes were identified by comparing all SGCs from the injured sample with those from the naive. The differential analysis was performed within each dataset to avoid adding batch effects and additional noise. For the Cell_SNI dataset, data were obtained 7 days and 14 days after nerve injury, while data for 3 days after injury was obtained for the Cell_crush dataset. Despite the differences in both time point and injury type, common differentially regulated genes were found to be enriched in related gene annotation groups (Figure 2D and Supplementary Excel Sheet: DE_analysis_metascape). For example, in both Cell_SNI at 14 days and Cell_crush at 3 days, genes were enriched in cholesterol biosynthesis. A closer look at the genes related to this ontology term reveals that the majority of them are downregulated in both datasets (Supplementary Figure 4 and Excel Sheet: DE_analysis_metascape).

To identify which specific regulated genes the Cell_SNI and Cell_crush datasets have in common the lists of differentially expressed genes were compared (Figure 2E). 18 genes were identified as common between Cell_SNI at 14 days and Cell_crush at 3 days – an enrichment that is 12x larger than expected by chance (as determined by hypergeometric probability calculations, assuming a total population of 10,000 genes as being expressed in SGCs). The common genes include several genes of within the cholesterol biosynthesis pathway: Idi1, Msmo1, Fdps, Fdft1 and Sqle (Table 3).

**Table 3:**
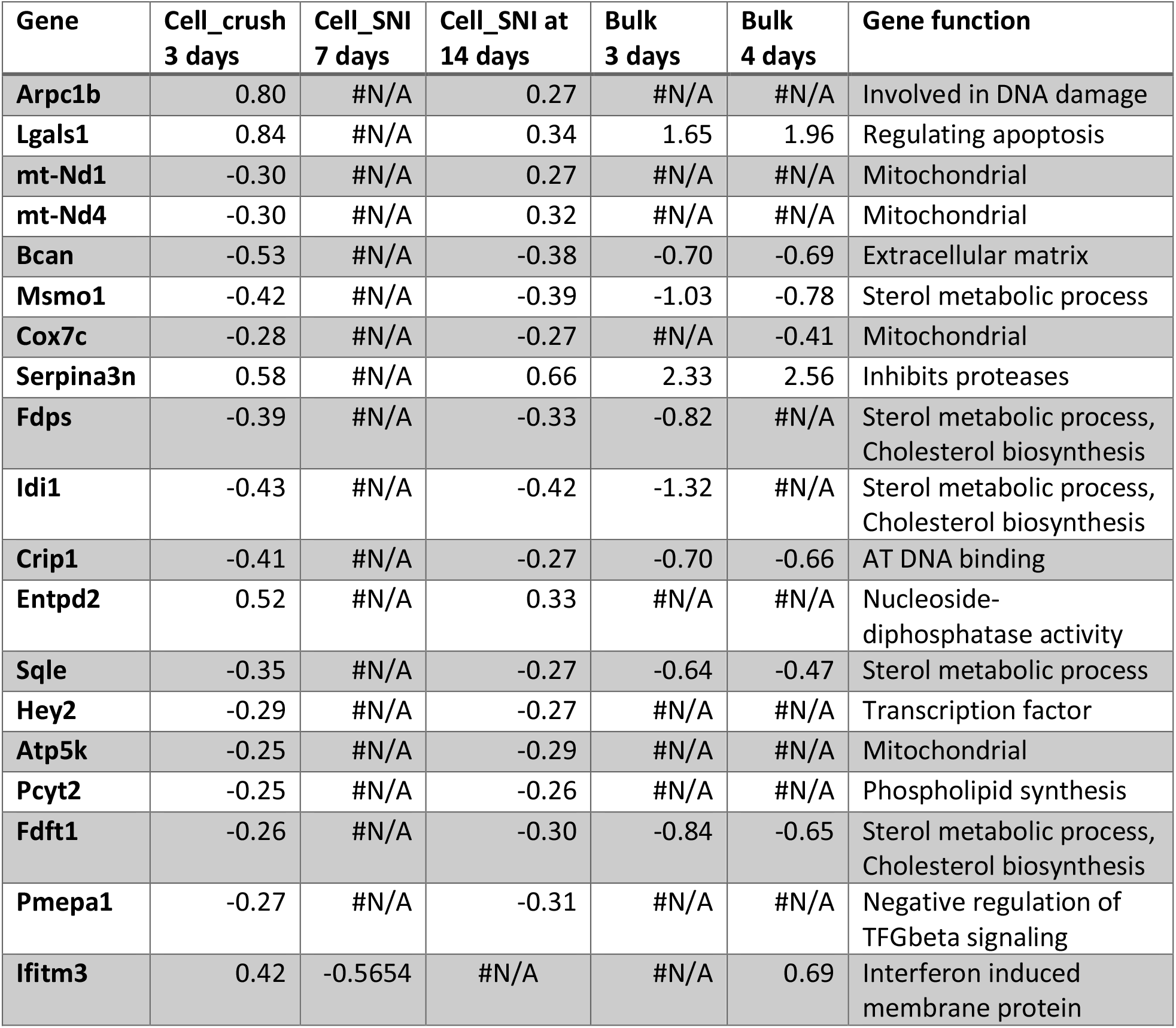
List of the 19 common regulated genes including log2 fold change and general gene function.

We have previously performed bulk RNAseq on sorted SGCs 3 and 14 days after transection of the sciatic nerve (Jager et al., 2020). To check whether scRNAseq and bulk RNAseq are in agreement, the 19 common genes identified between the Cell_SNI and Cell_crush datasets were compared to the gene expression in the bulk dataset (Figure 2F). Of these 19 genes, 11 genes are also significantly regulated in the bulk dataset and in the same up/down direction (Figure 2F and Table 3), confirming the regulation of the genes involved in cholesterol biosynthesis (Idi1, Msmo1, Fdps, Fdft1 and Sqle).

### Regulation of known SGC markers

The list of common regulated genes (Table 3) includes several that have yet to be investigated in the context of SGC function. Surprisingly, the list did not include genes that have previously been reported to be regulated at protein or gene level such as Connexin43 (*Gja1*), GFAP (*Gfap*) or Hmgcs1 (*Hmgcs1*) (Ohara, Vit, Bhargava, & Jasmin, 2008; F. Wang et al., 2016; Komiya et al., 2018). Therefore, these genes were further examined in the datasets (2x scRNAseq, 1x bulk RNAseq). Connexin43 has been shown to be increased at protein level in SGCs after nerve injury (Ohara et al., 2008; Komiya et al., 2018). Counterintuitively, a downregulation of *Gja1* at the mRNA level in the bulk RNAseq were observed while no regulation of *Gja1* in the scRNAseq datasets were detected (Table 4).

**Table 4:** Analysis of differential expression of Gja1, Gfap and Hmgcs1 in SGCs in various datasets. % = % of SGCs expressing Gja1 (gene for Connexin43), Gfap and Hmgcs1. FPKM = Fragment per kilobase of transcript per million mapped reads.

| Dataset and time point | % | FPKM | Log2 foldchange | Adj P-value |
| --- | --- | --- | --- | --- |
| **Gja1** |  |  |  |  |
| Cell_SNI 7 days | 47 | N/A | -0.05 | N/A |
| Cell_SNI 14 days | 40 | N/A | -0.24 | N/A |
| Cell_crush 3 days | 31 | N/A | -0.20 | N/A |
| Bulk 3 days | N/A | 88 | -0.58 | 0.02 |
| Bulk 14 days | N/A | 100 | -0.46 | 0.001 |
| **Gfap** |  |  |  |  |
| Cell_SNI 7 days | 3 | N/A | 0.1 | N/A |
| Cell_SNI 14 days | 7 | N/A | 0.24 | N/A |
| Cell_crush 3 days | 3.7 | N/A | 0.26 | N/A |
| Bulk 3 days | N/A | 0.33 | 1.8 | N/A |
| Bulk 14 days | N/A | 0.41 | 2.2 | N/A |
| **Hmgcs1** |  |  |  |  |
| Cell_SNI 7 days | 71 | N/A | -0.1 | N/A |
| Cell_SNI 14 days | 72 | N/A | -0.24 | N/A |
| Cell_crush 3 days | 66 | N/A | -0.43 | $1.2 * 10^{-29}$ |
| Bulk 3 days | N/A | 217 | -0.93 | $1.9 * 10^{-14}$ |
| Bulk 14 days | N/A | 269 | -0.69 | $4 * 10^{-9}$ |

Increased expression of GFAP protein is often used as a marker for SGC reactivity by immunohistochemical analysis (Ohara et al., 2008; M Hanani, Blum, Liu, Peng, & Liang, 2014; Blum, Procacci, Conte, Sartori, & Hanani, 2017; Komiya et al., 2018). In bulk RNAseq, *Gfap* gene expression was not detected above threshold (FPKM>1), as we previously described (Jager et al., 2020). In accordance with this, expression of *Gfap* in the scRNAseq datasets (Cell_SNI and Cell_crush) were only detected in 3 – 7% of SGCs, which is below our defined threshold (see methods). Furthermore, we did not observe differential regulation of *Gfap* above threshold in either dataset (Table 4). Whether this result reflects strain or species variation is discussed elsewhere (Mohr, Pallesen, Richner, & Vaegter, 2021).

Finally, the cholesterol synthesis pathway enzyme Hmgcs1 has been shown to be downregulated in SGCs after nerve injury (F. Wang et al., 2016). In our datasets, significant transcriptional downregulation of *Hmgcs1* in the Cell_crush dataset and the bulk RNAseq after sciatic nerve ligation (Table 4) was detected. We did not confirm *Hmgcs1* downregulation in the Cell_SNI dataset, however downregulation of other genes involved in the biosynthesis of cholesterol were observed, supporting injury-induced regulation of the cholesterol synthesis pathway in SGCs (Supplementary Figure 4 and Supplementary Excel Sheet: DE_analysis_metascape).

### Transcriptional response in cultured glia cells

SGCs have on several occasions been investigated using *in vitro* cultures from either pups or adult rodents (Fornaro, Sharthiya, & Tiwari, 2018; Leisengang et al., 2018; Vinterhøj, Stensballe, Duroux, & Gazerani, 2019; X. bin Wang et al., 2019). However, reports of loss of marker protein expression upon disconnection from their associated neuron (Belzer et al., 2010) as well as regression to a transcriptional profile closely identical to that of Schwann cell precursor-like state (George et al., 2018) complicate meaningful translational interpretations to the *in vivo* condition. Here scRNAseq was performed on primary cultures of mouse DRGs (Cell_culture, GSE188971) to compare the transcriptional profiles of such cultured SGCs to that of acutely isolated naïve and injured SGCs of the Cell_SNI dataset. When performing the initial cluster analysis of the Cell_culture dataset, 4 different clusters of cells were identified: neurons, macrophages, fibroblasts and glial cells (Figure 3A), with glial cells constituting the vast majority (88%). The glia cell cluster was explored in the attempt to subdivide further by relying merely on the expression of traditional Schwann cell and SGC markers (Figure 3B). However, the SGC markers *Fabp7* and *Kcnj10* (Kir4.1) showed no clear SGC clustering, and Schwann cell markers were even more widely dispersed, indicating that glial cells change their gene expression profiles extensively *in vitro*.

**Figure 3:**
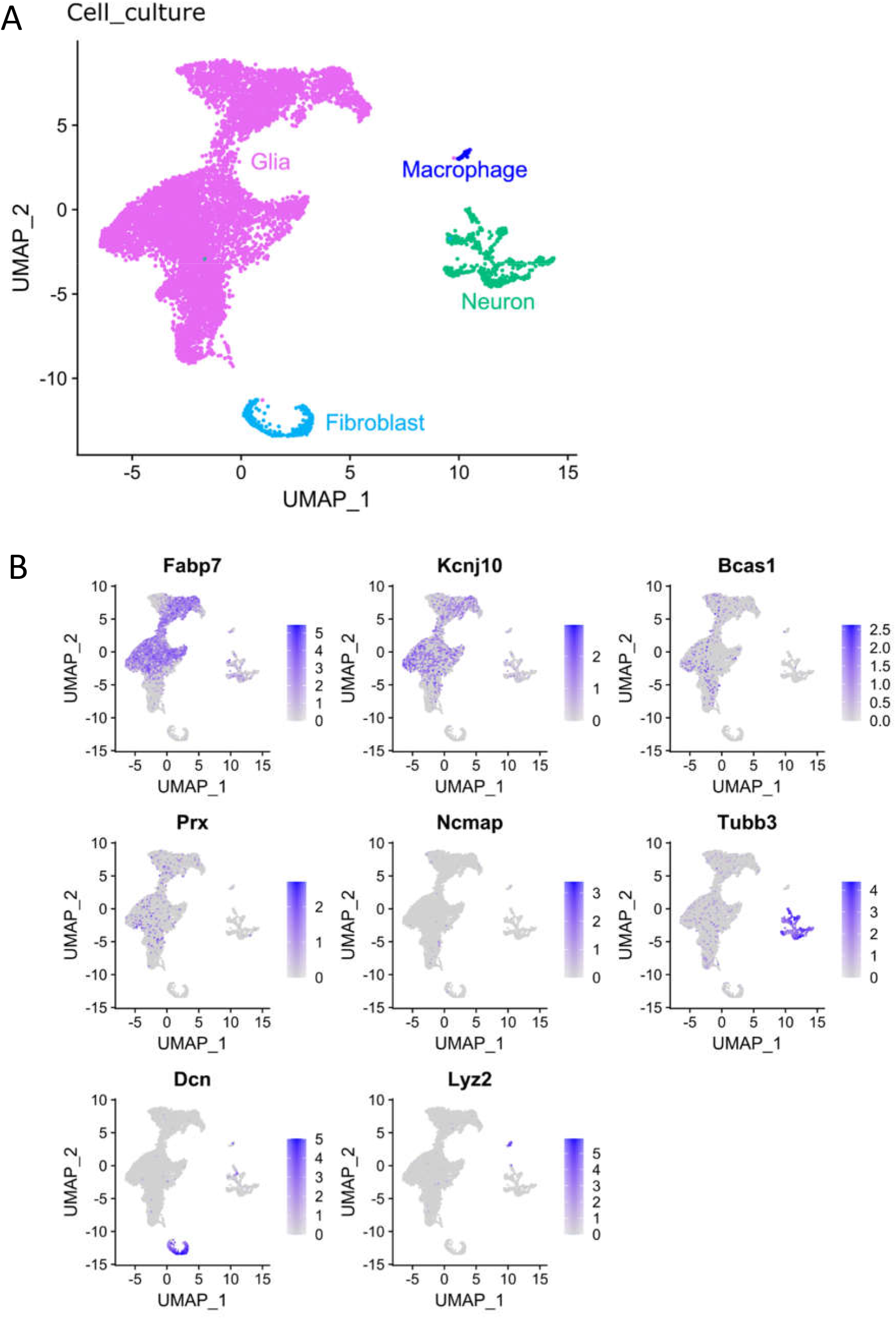
The cell_culture dataset contains glial cells, fibroblasts, neurons, and macrophages. A) UMAP of the identified cell types. B) Expression of markers for SGCs (Fabp7 and Kcnj10), Schwann cells (Bcas1, Prx, Ncmap), neurons (Tubb3), fibroblasts (Dcn) and macrophages (Lyz2).

To improve annotations and investigate translational variations of cultured SGCs relative to their *in vivo* state, the Cell_culture dataset was integrated with the Cell_SNI dataset to enable joint analyses (Figures 4A and 4B). Cell culture glia cells clustered together with acutely isolated SGCs and Schwann cells (Figure 4B). A projection of the integrated annotation back onto the Cell_culture dataset pre-integration is illustrated in Figure 4C and shows that the glia cluster (Figure 3A) indeed contains many different cell types. The joint analysis also reveals a distinct glial cell cluster (“*In vitro* glia”), selectively present in the Cell_culture dataset (Figure 4D). These cells are enriched for SGC marker genes such as *Fabp7* and *Kcnj10*, and for genes involved in cell proliferation, such as *Top2a* (DNA topoisomerase II alpha) and *Mki67* (marker of proliferation Ki-67) (Figure 4E). It has previously been suggested by George *et al*. that peripheral glia cells regress back to a Schwann cell precursor (SCP) phenotype when cultured (George et al., 2018). To investigate if this could be the fate of the “*in vitro* glia” cluster, the joint dataset was compared to a dataset containing cells from the developing peripheral nervous system at E12.5 (Furlan et al., 2017) with the R package SingleR (Aran et al., 2019). The analysis shows that the “*in vitro* glia” cluster does indeed resemble SCPs (Figure 4F) supporting the hypothesis that peripheral glia regress into a SCP phenotype in culture. As expected, the neuronal cluster has similarities with sympathoblasts which develop into sympathetic neurons and chromaffin cells which are neuroendocrine cells (Figure 4F).

**Figure 4:**
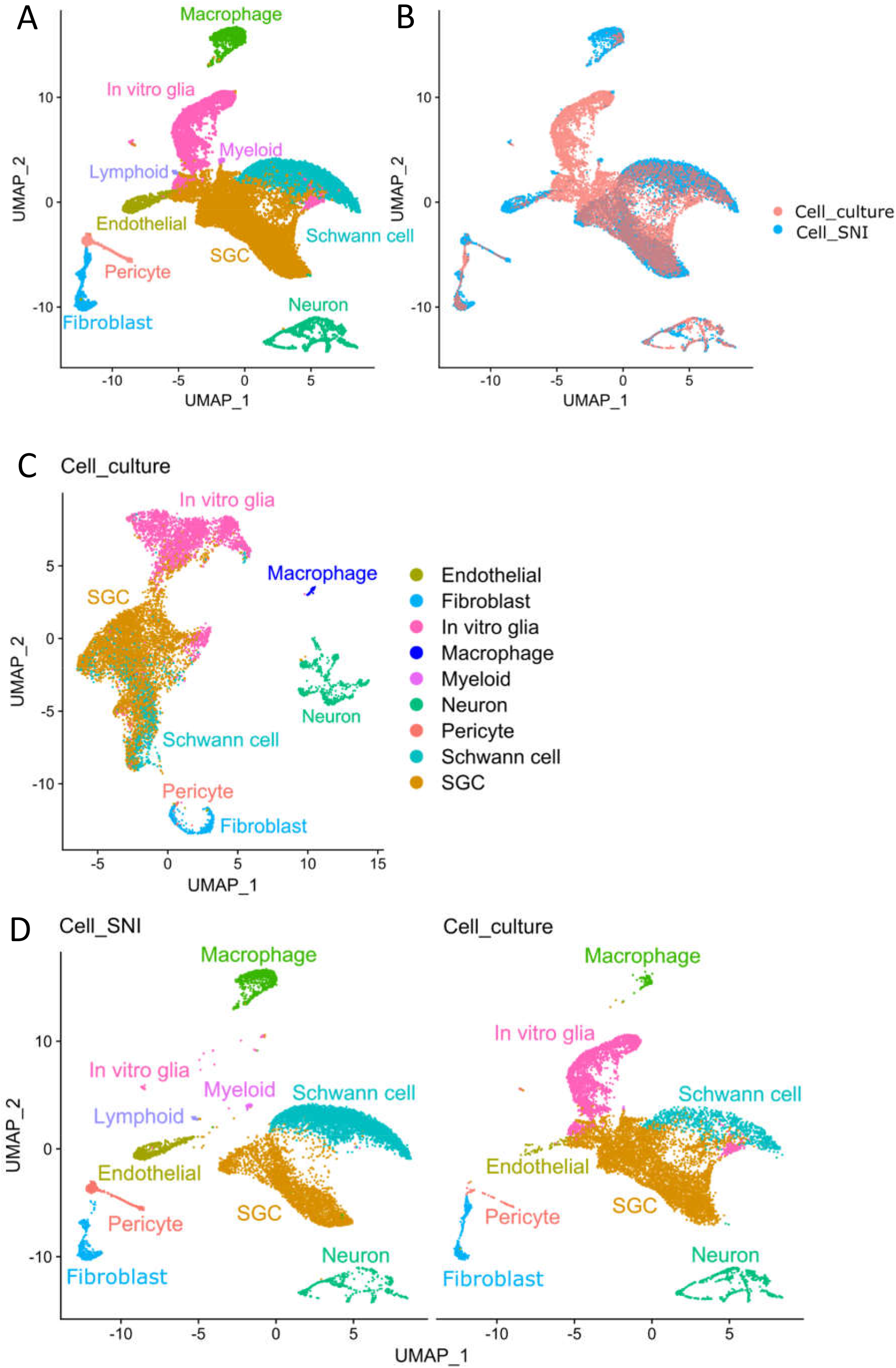
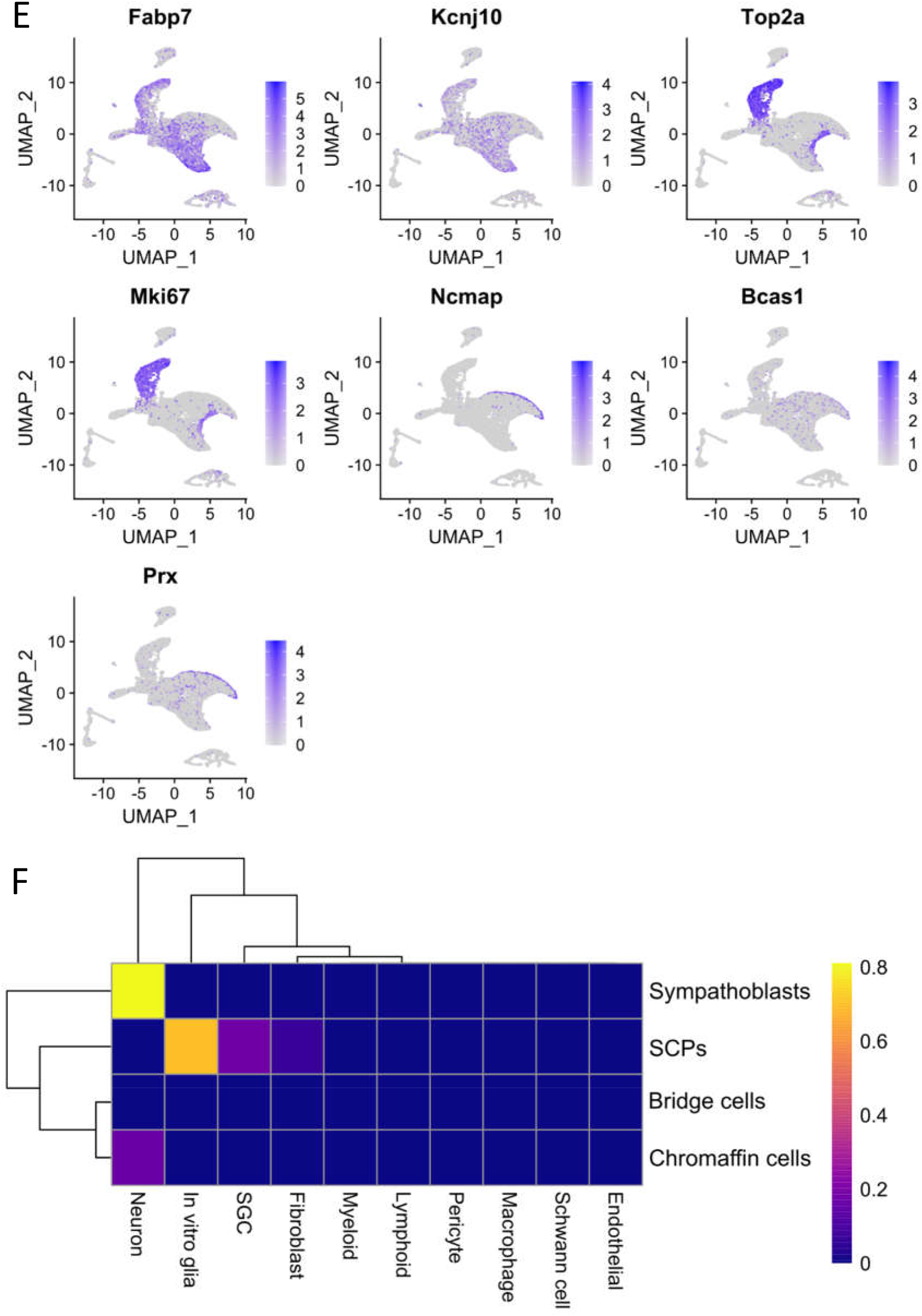
Glia cells change in culture. A) UMAP of joint analysis with annotation of cell types. B) UMAP of joint analysis coloured based on dataset. C) UMAP of Cell_culture dataset with annotation from joint analysis. D) UMAPs of individual datasets with annotation of cell types identified from the joint analysis. E) Expression of markers for SGCs (Fabp7 and Kcnj10), cell proliferation (Top2a and Mki67) and Schwann cells (Ncmap, Bcas1 and Prx). F) Heat map showing the result of the SingleR analysis which compared the gene expression in the joint analysis with cells in the developing peripheral nervous system. SCP = Schwann cell precursor.

### SGCs change toward a precursor phenotype *in vitro*

Finally, the joint analysis was used to identify differences between the SGCs originating from the Cell_culture or the Cell_SNI dataset (orange cluster in Figure 4A). To increase the resolution for the SGC cluster, it was subset and re-clustered. This showed that a significant number of the cells from the Cell_culture condition cluster separately (Figure 5A and B), suggesting that their transcriptional profile diverges significantly from those of Cell_SNI SGCs. This was particularly the case for cells in subclusters 2 and 3 (Figure 5B).

**Figure 5:**
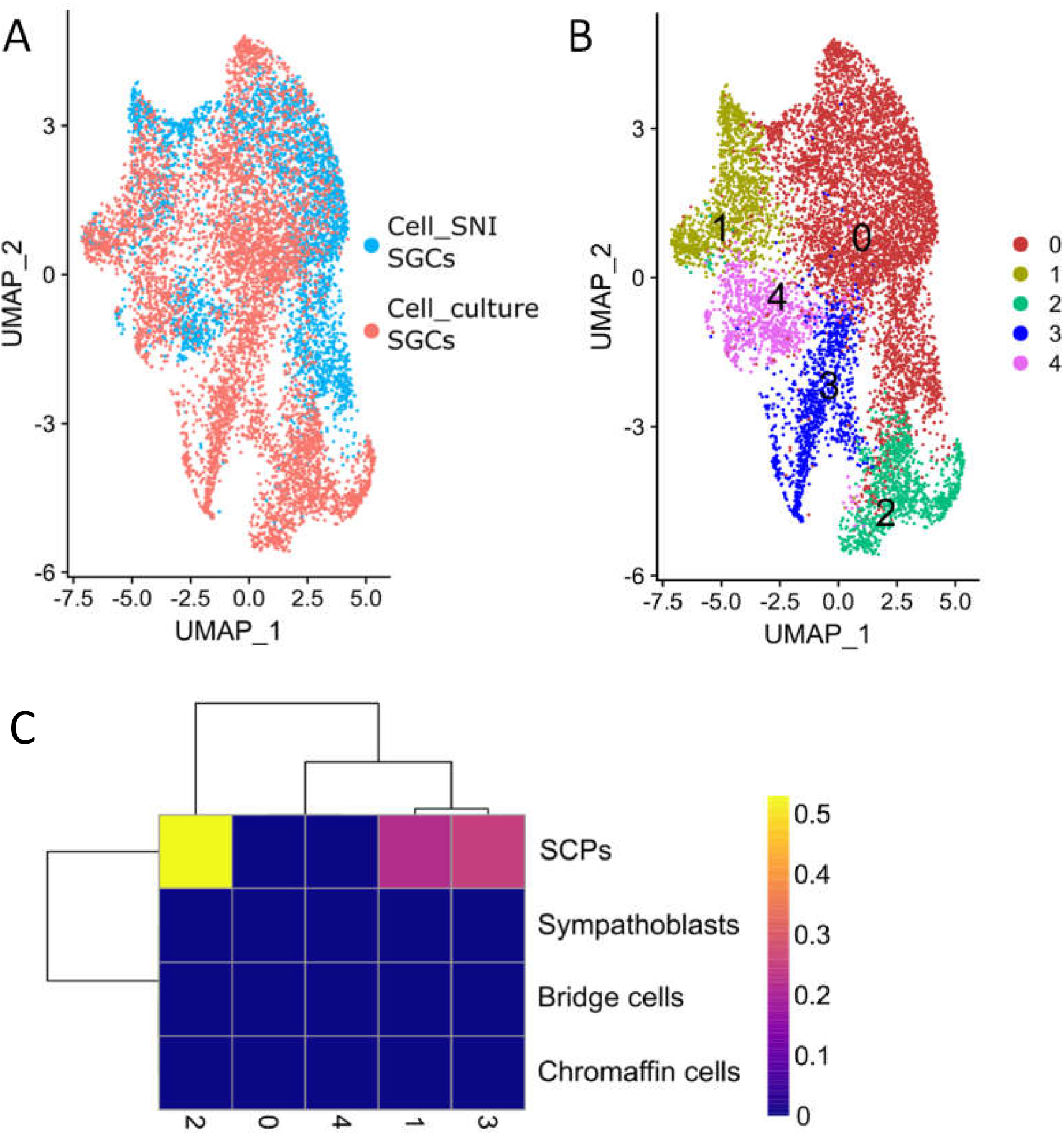
Some SGC change towards a precursor phenotype in vitro. A) UMAP of SGC cluster from the joint analysis coloured by dataset. B) UMAP of SGC cluster from joint analysis with new cluster analysis. C) Heat map showing the result of the SingleR analysis which compared the gene expression in the SGC clusters with cells in the developing peripheral nervous system. SCP = Schwann cell precursor.

To investigate whether these culture-induced changes also point to a regression towards a SCP phenotype, the 5 SGC clusters were compared to the cell types in the developing nervous system (Furlan et al., 2017) with SingleR (Aran et al., 2019). The results revealed that clusters 2 and 3 resemble SCPs (Figure 5C), raising the possibility that a proportion of SGCs in culture revert to a mutual precursor phenotype.

In conclusion, our analyses of the joint dataset revealed that only part of the SGCs in culture resembles *in vivo* SGCs (cluster 0, 1 and 4 of Figure 5B) and identified two changes in culture: the appearance of a culture specific cell type (the “*In vitro* glia” of Figure 4C), which express SGC markers, proliferation genes and resembles SCPs and a proportion of the SGCs with a SCP phenotype (SGC clusters 2 & 3 of Figure 5B).

## Discussion

In the last few years, single nucleus and single cell RNAseq datasets have been published to investigate the injury response of DRG cells, with a particular focus on neurons (Renthal et al., 2020; K. Wang et al., 2021) and SGCs (Avraham et al., 2020). With this study, we adopted a meta-scientific approach to summarise specifically the over-arching conclusions that can be drawn from these data on how SGCs behave after nerve injury. From the 4 datasets we considered for inclusion, two (Renthal et al., 2020; K. Wang et al., 2021) were excluded due to high levels of neuronal contamination in the differential expression analysis. The reasons for this contamination are not clear. As all the investigated non-neuronal cell types and not only the SGCs have the ‘canonical’ neuronal response, we find it unlikely that it should be due to insufficient disruption of the SGC-neuron units. Instead, we believe that it may be related to the magnitude of transcriptional regulation in neurons, which dwarfs that of all other DRG cell types following nerve injury. This greater response can be a source of cross-contamination, if neuronal mRNA is present in the cellular mixture before droplet separation. In the case of Renthal *et al*., significant amounts of cytosolic mRNA would have been released during the isolation of nuclei just prior to their single-nucleus RNAseq. In the case of Wang *et al*., their neuronal enrichment step result in more neurons being sequenced than in the SGC-focused datasets. We speculate that this would also have been accompanied by a proportional increase in the number of dead neurons (i.e. free neuronal mRNA) in the starting cell mixture.

Analysing the two remaining datasets, we identified a common SGCs transcriptional injury response, with downregulation of genes annotated to cholesterol biosynthesis. This finding is in line with protein data published by Wang *et al*. (F. Wang et al., 2016), who reported downregulation of the cholesterol pathway protein Hmgcs1 in rat DRG after spinal nerve ligation. Little is known about the possible functional consequences of this potential change in cholesterol metabolism. After nerve injury, it has been shown that SGCs increase their cell membrane surface area (E Pannese et al., 2003). It seems counter-intuitive that there can both be a downregulation of cholesterol production and an increased membrane production, considering mammalian plasma membranes consist of app. 30% cholesterol (Krause, Daly, Almeida, & Regen, 2014). One might wonder whether SGCs change how they obtain their cholesterol after nerve injury. Since they express general cholesterol receptors, like LDLR and VLDLR, they would be capable of taking up cholesterol from the extracellular space, where it might be released from activated macrophages. Macrophages are known for their high cholesterol production, and we and others have shown that they increase in number and migrate into the SGC-neuron unit after injury (Lu & Richardson, 1993; Dubový, Tucková, Jancálek, Svíženská, & Klusáková, 2007; Vega-Avelaira, Geranton, & Fitzgerald, 2009; Jager et al., 2020; Iwai et al., 2021). At present, however, this remains speculation until more functional data can be obtained.

When performing sciatic nerve injuries on mice, not all neurons in the corresponding DRG (L3-L5) will be injured (Rigaud et al., 2008). Consequently, we expect not all SGCs in the injured samples to have an injury response. We were therefore surprised to see that SGCs did not cluster in two groups based on whether they surrounded injured neurons or not. We speculate that the transcriptional response is too subtle to allow for sub-clustering of the SGCs into injured and uninjured cells, at least amongst the transcripts we were able to capture with droplet-based methods and the 4581 SGCs (2016 SGCs from Cell_crush and 2565 SGCs from Cell_SNI) analysed here.

Beyond the examination of acutely isolated SGCs, we also studied those that had been cultured for 3 days. Our results indicate that the gene expression profile of cultured peripheral glial cells changes significantly *in vitro*. We found that an entirely new population emerges upon culturing which we labelled “*in vitro* glia”. It is characterized by expression of genes related to proliferation, expression of SGCs markers and a resemblance to Schwann cell precursors. In addition to these “*in vitro* glia”, we also found that a proportion of cells within the “more physiological” SGC cluster in culture, change into a Schwann cell precursor-like state. This is in line with work from George *et al*., who showed that long-term cultured SGCs have a similar transcriptional profile to that of long-term cultured Schwann cells (George et al., 2018). Our cultured cells were derived from 2-week old mice, where the maturation of promyelinating Schwann cells to myelinating Schwann cells is still in process (Monk, Feltri, & Taveggia, 2015). We therefore cannot exclude that this developmental timeline for Schwann cells had an impact in our results.

The proliferation profile seen in the “*in vitro* glia” is absent in acutely isolated SGCs. Specifically, at least transcriptionally, we did not find any evidence to suggest that adult SGCs cells are proliferating after nerve injury *in vivo*. Reports to the contrary (Friede & Johnstone, 1967; Lu & Richardson, 1991; J. P. Vit et al., 2006; Donegan, Kernisant, Cua, Jasmin, & Ohara, 2013) are confounded by the fact that they stained only for proliferation markers and attempted to identify SGCs by their position rather than by antibody staining. Especially after nerve injury, when macrophages closely approach SGC-neuron units, this intimate position of macrophages relative to the neuronal soma may easily be misinterpreted as SGCs when omitting detection of cellular markers (Jager et al., 2020). During development, SGCs and other cells do proliferate in the DRG, but this process has been shown to terminate around birth (Lawson & Biscoe, 1979; George et al., 2018).

Like all single-cell studies, our analysis had limitations. Importantly, most current scRNA-seq experiments, including those presented here, rely on droplet-based technologies that are only able to detect a fraction of transcripts present in a given cell (∼30%) (10X Genomics, 2021). In the Cell_SNI dataset, we analysed 2565 SGCs, suggesting that across all SGCs, we are likely to have a good representation of the genes detectable in SGCs. Indeed, when we compiled all single cell transcripts to generate a pseudo-bulk profile, we found comparable expression to our own prior bulk sequencing results of sorted SGCs (see Supplementary Excel Sheet: SGC_gene_expression). Nevertheless, with either method, we may have missed very lowly expressed transcripts, like adhesion GPCRs (due to the number of cells analysed here, and the read depth used in (Jager et al., 2020)).

Our differential expression analyses were generally rather variable – as indicated by the low number of commonly regulated genes identified across datasets. One possible explanation is the difference in time points and injury types. For instance, nerve crush is a regenerating model, while SNI is a chronic model causing persistent pain. Another likely cause for the observed variability is that we were limited to performing the differential expression analyses on a cell-by-cell basis, an approach which lacks power and gives rise to a higher frequency of false positives. If we had had more biological replicates, we could have performed a pseudo-bulk analysis which might have shed further light on the common responses of SGCs to different nerve injuries (Crowell et al., 2020; Squair et al., 2021).

Finally, as with all scRNAseq studies, we have to assume that the use of enzymes and/or mechanical forces to obtain a single cell suspension prior to sequencing will, in itself, alter expression of some genes (van den Brink et al., 2017).

In conclusion, we found that SGCs share a common response following nerve crush and ligation, which includes regulation of genes involved in cholesterol biosynthesis. We also found that peripheral glial cells in culture change significantly, with many starting to resemble Schwann cell precursors. Our *in vitro* observations were in accordance with previous studies (Belzer et al., 2010; George et al., 2018) and emphasize how studies using SGC in a dish need to be approached and interpreted with caution.

## Supporting information

Supplementary SGC_gene_expression

Supplementary Wang et al 7d

Supplementary DE_analysis_metascape

Supplementary Renthal et al 7d

Supplementary_notebook

## Acknowledgements

We would like to thank Dr William Renthal for generously sharing their data in an easily accessible format ahead of their formal publication (Renthal et al., 2020). We would like to thank Prof Patrizia Rizzu and her research team at the DZNE in Tübingen for their contribution to the 10X sequencing of the Cell_Culture dataset.

## Data availability

A significant portion of the information available in the single cell datasets used in this study is not easily accessible to most wet lab researchers. To mitigate this, we have uploaded the scRNAseq studies used in this analysis to the single cell web-based portal at Broad Institute www.singlecell.broadinstitute.org/single_cell/study/SCP1539/.

The fastq files, count matrices and Seurat objects can be found on the Gene Expression Omnibus repository under GSE174430 for the Cell_SNI dataset, GSE139103 for the Cell_Crush dataset and GSE188971 for the Cell_culture dataset. The code to make the R based figures can be found in the Supplementary notebook.

