## Supplementary figures for "Comparative transcriptional analysis of the satellite glial cell injury response"

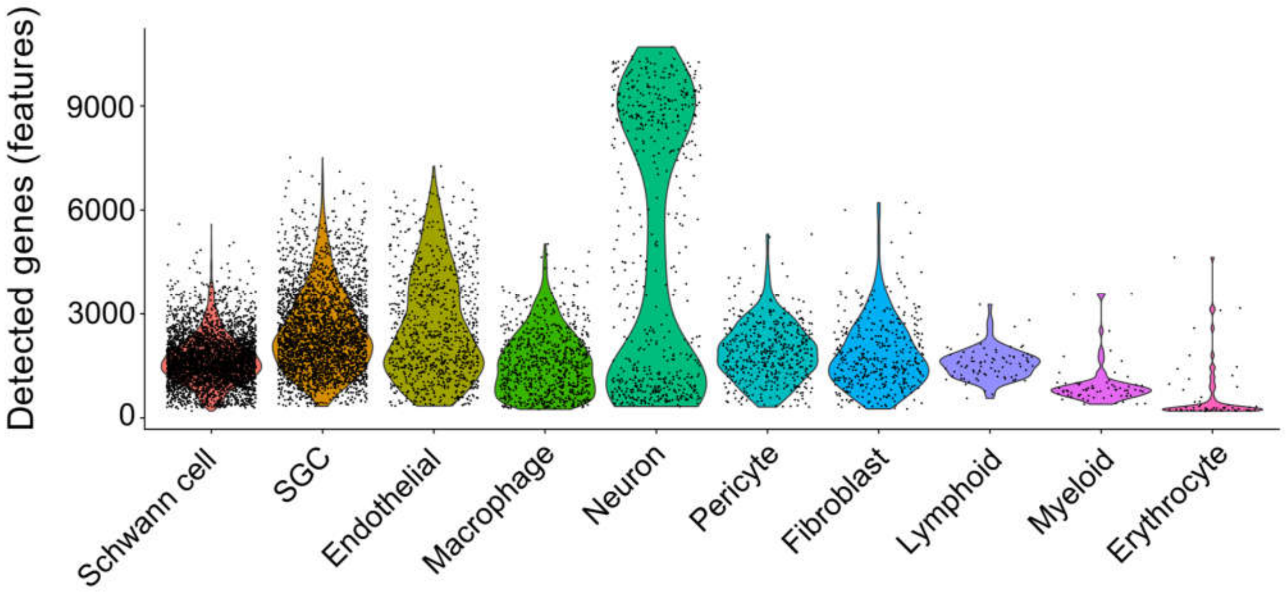


*Supp. figure 1 Plot showing the cell types from the Cell_SNI dataset on the x-axis and the number of detected genes on the y-axis*

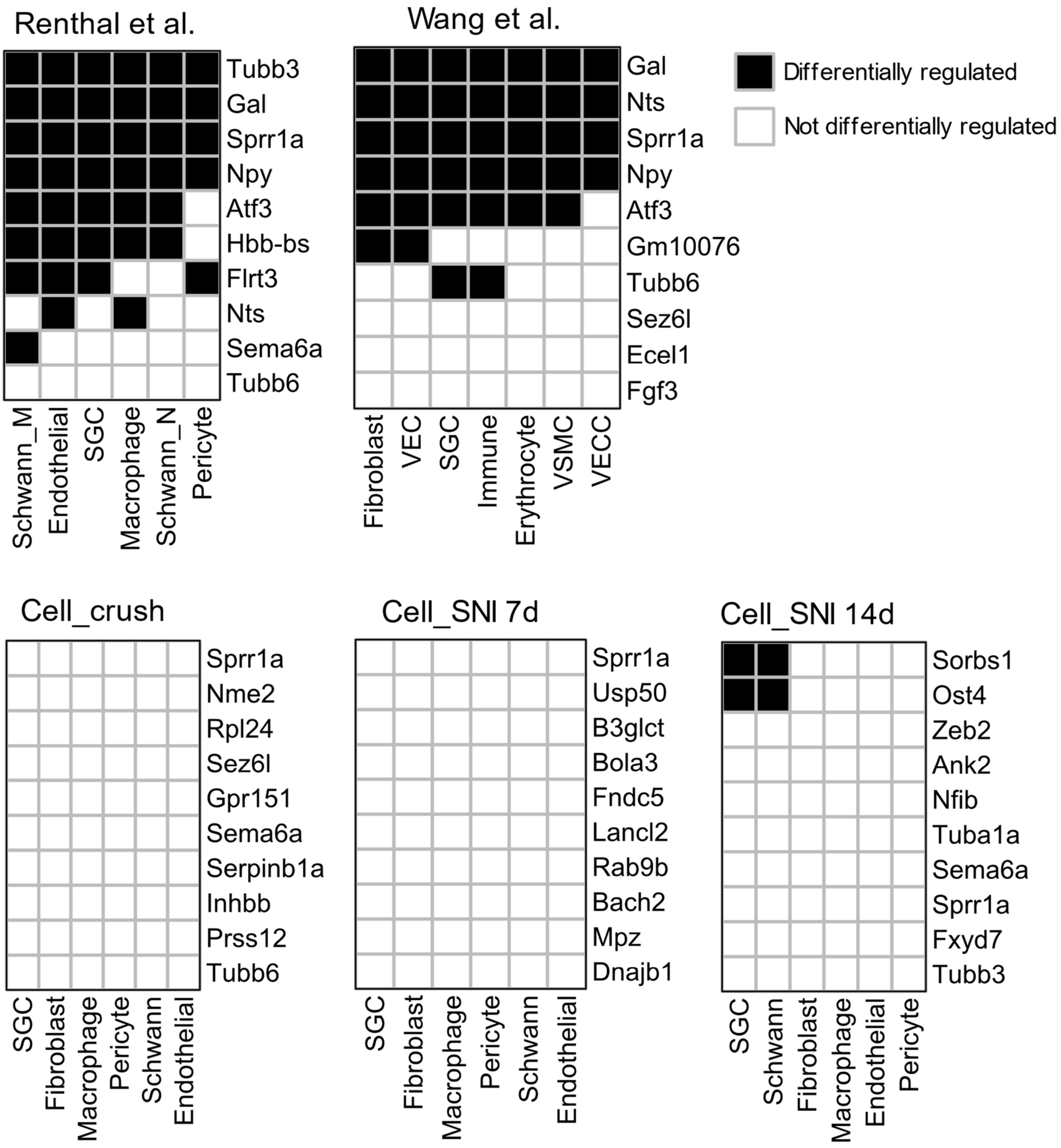


*Supp. figure 2: Heatmaps with the top 10 regulated neuronal genes plotted against the non-neuronal cell type clusters identified in each dataset. Black denotes that a gene is differentially regulated in the corresponding non-neuronal cluster and white that it is not regulated. SGC = satellite glial cells, VEC = vascular endothelial cells, VSMC = vascular smooth muscle cells and VECC = vascular endothelial capillary cells.*

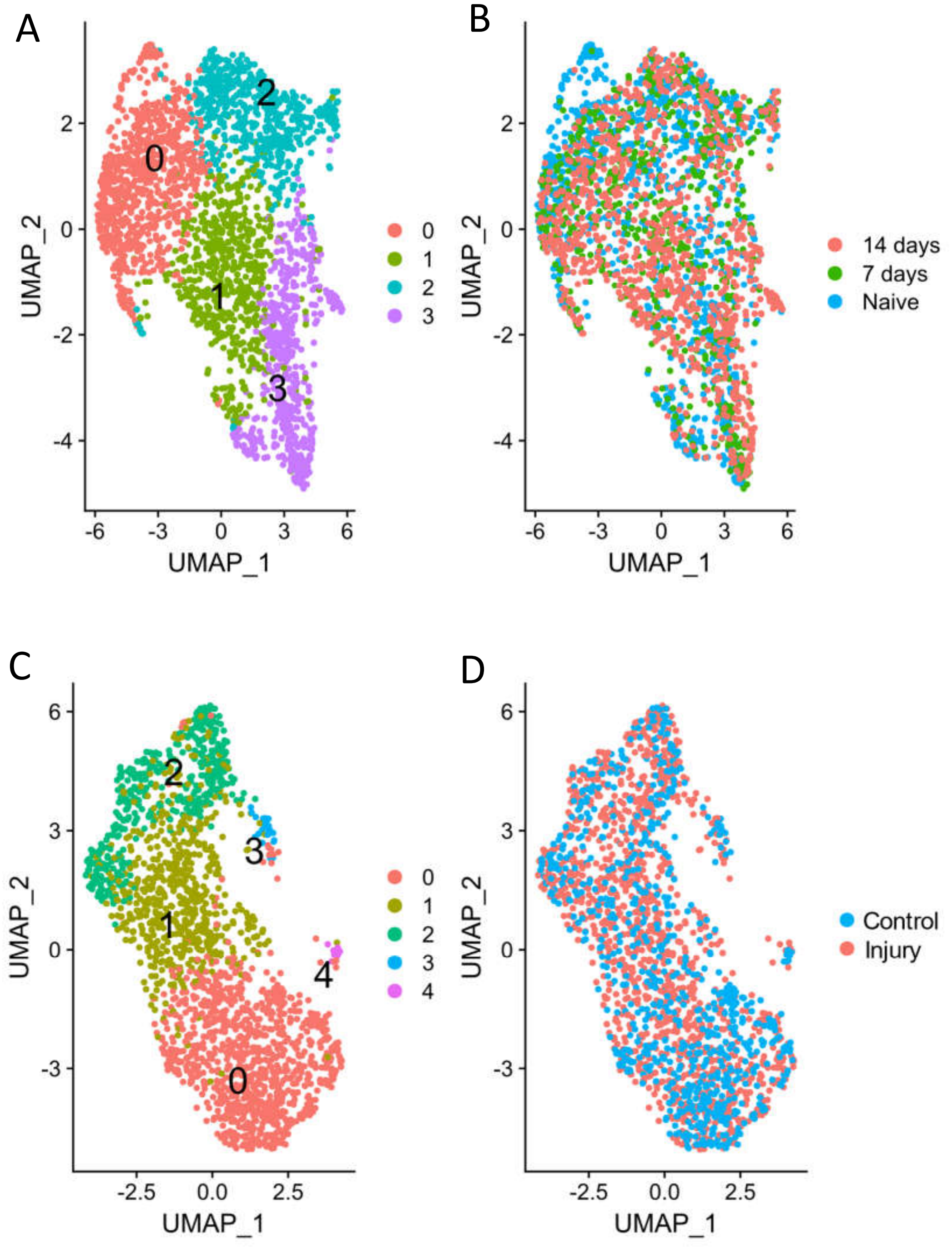


*Supp. figure 3: SGCs clusters in Cell_SNI and Cell_crush datasets. A-B) UMAPS of the SGCs from the Cell_SNI dataset. A) UMAP coloured based on clustering B) UMAP coloured based on injury condition. No cluster contains only injured SGCs. C-D) UMAPS of the SGCs from the Cell_crush dataset. C) UMAP coloured based on clustering. D) UMAP coloured based on injury condition. No cluster contains only injured SGCs.*

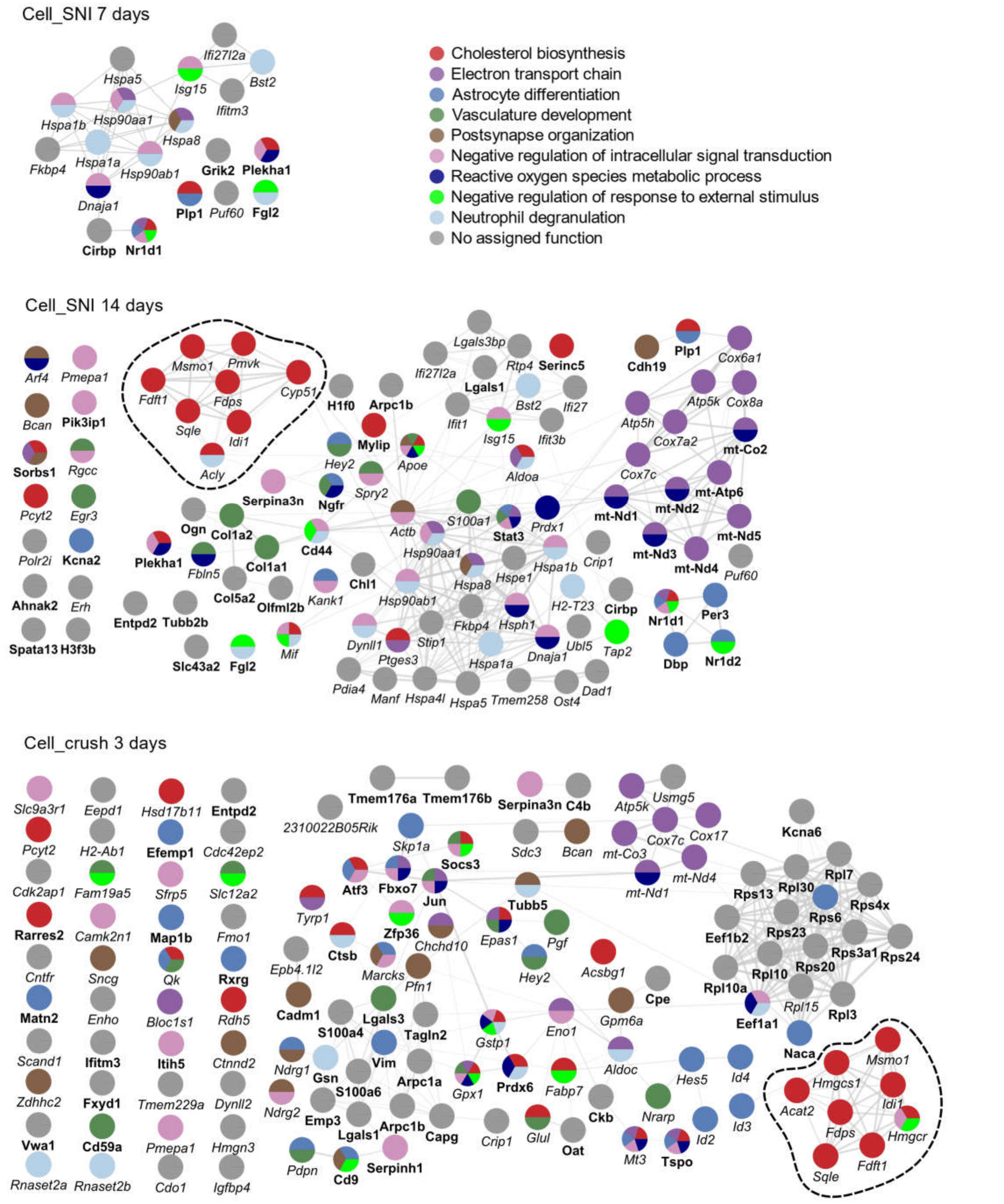


*Supp. Figure 4: STRING network of the regulated genes in the datasets. The Cell_SNI dataset is spilt based on timepoints. The genes are colour coded based on their annotation related to the 9 terms that both datasets had in common in Figure 2D. Gene names in bold indicate that the gene was upregulated after injury while italic gene names indicate that the gene was downregulated. The dashed line highlights the downregulated genes involved in the cholesterol biosynthesis pathway.*
