## Supplementary_notebook for "Comparative transcriptional analysis of the satellite glial cell injury response": Supplementary_notebook.nb.html

Notebook\_for\_figures\_made\_in\_R


Code 

- Show All Code
- Hide All Code
- Download Rmd

### Notebook\_for\_figures\_made\_in\_R

This notebook contains the code used to make the R based figures. Most of the analysis is done with Seurat. ScMap is used to project one dataset on to another and SingleR is used to compare gene expression profiles across datasets.


```
#The following packages are needed. Before you can open the packages with the library() function you need to download them. If in doubt search for download instructions for each package on google. 
library(Seurat)
library(scmap)
library(cowplot)
library(patchwork)
library(dplyr)
library(scater)
library(ggplot2)
library(scales)
library(SingleR)
library(Matrix)
library(SingleCellExperiment)
library(pheatmap)
library(viridis)
```


First we load the data for the Cell\_SNI dataset. This file can be downloaded from GEO accession number GSE174430 under supplementary files.


```
Cell_SNI <- readRDS("C:/Users/your/path/Cell_SNI_data.rds") #Adjust the path to fit your system
```


To get FIGURE 1A with the Cell\_SNI dataset:


```
DimPlot(Cell_SNI,reduction = "umap", label = TRUE, repel = TRUE) + NoLegend()
```


Next we need to load the data from Cell\_crush.This is a bit more complicated because we have to download the count matrix from the original publication and re-analyse it to annotate the cell types.

Download the GSE139103\_RAW.tar file from GSE139103 under supplementary files. Extract the files on your PC. This should give you 4 folders named: GSM4131028\_CON, GSM4131029\_INJ, GSM4131030\_CON2 and GSM4131031\_INJ2. The folders contain the output files from Cell Ranger that we need to make a Seurat object.

First I read the files in and next I merge them so I get one seurat object with all the crush data


```
con1_data <- Read10X("C:/Users/your/path/GSM4131028_CON") #Adjust the path to fit your system
con1 <- CreateSeuratObject(con1_data, "Con1")
con1
```


```
inj1_data <- Read10X("C:/Users/your/path/GSM4131029_INJ") #Adjust the path to fit your system
inj1 <- CreateSeuratObject(inj1_data, "Inj1")
inj1
```


```
con2_data <- Read10X("C:/Users/your/path/GSM4131030_CON2") #Adjust the path to fit your system
con2 <- CreateSeuratObject(con2_data, "Con2")
con2
```


```
inj2_data <- Read10X("C:/Users/your/path/GSM4131031_INJ2") #Adjust the path to fit your system
inj2 <- CreateSeuratObject(inj2_data, "Inj2")
inj2
```


```
Cell_crush <- merge(inj1, y = c(con1, inj2, con2), project = "Avraham_crush")
Cell_crush
```


Next we make a new meta data column called percen.mt to check how much mitochondrial message we got.


```
Cell_crush[["percent.mt"]] <- PercentageFeatureSet(Cell_crush, pattern = "^mt-")
```


Next we check the quality of the data:


```
VlnPlot(Cell_crush, features = c("nFeature_RNA", "nCount_RNA", "percent.mt"), ncol = 3)
```


I filter out cell with less than 200 detected genes and with more than 15% mitochondrial genes.


```
Cell_crush <- subset(Cell_crush, subset = nFeature_RNA > 200 & percent.mt < 15) #Here is the filtering done
VlnPlot(Cell_crush, features = c("nFeature_RNA", "nCount_RNA", "percent.mt"), ncol = 3) #Here I display the data again
```


Next I need to integrate the samples to mitigate batch effects:


```
# split the dataset into a list of two seurat objects
ifnb.list <- SplitObject(Cell_crush, split.by = "orig.ident")

# normalize and identify variable features for each dataset independently
ifnb.list <- lapply(X = ifnb.list, FUN = function(x) {
    x <- NormalizeData(x)
    x <- FindVariableFeatures(x, selection.method = "vst", nfeatures = 2000)
})

# select features that are repeatedly variable across datasets for integration
features <- SelectIntegrationFeatures(object.list = ifnb.list)
```


Perform integration


```
crush_anchors <- FindIntegrationAnchors(object.list = ifnb.list, anchor.features = features, dim=1:20)
# this command creates an 'integrated' data assay
Cell_crush <- IntegrateData(anchorset = crush_anchors, dim=1:20)
```


```
# specify that we will perform downstream analysis on the corrected data note that the original
# unmodified data still resides in the 'RNA' assay
DefaultAssay(Cell_crush) <- "integrated"

# Run the standard workflow for visualization and clustering
Cell_crush <- ScaleData(Cell_crush, verbose = FALSE)
Cell_crush <- RunPCA(Cell_crush, npcs = 30, verbose = FALSE)
Cell_crush <- RunUMAP(Cell_crush, reduction = "pca", dims = 1:20)
Cell_crush <- FindNeighbors(Cell_crush, reduction = "pca", dims = 1:20)
Cell_crush <- FindClusters(Cell_crush, resolution = 0.08)
```


```
# Visualization
DimPlot(Cell_crush, reduction = "umap", group.by = "orig.ident")
```


```
DimPlot(Cell_crush, reduction = "umap", label = TRUE, repel = TRUE)
```


The UMAP generated here might look a bit different to the once in the paper. This is because UMAPs have a random component.

Now we need to find the cell type markers to annotate the clusters in the UMAP above.


```
DefaultAssay(Cell_crush) <- "RNA" #I switch to the unintegrated data to find cell type markers

# find markers for every cluster compared to all remaining cells, report only the positive ones
Cell_crush_markers <- FindAllMarkers(Cell_crush, only.pos = TRUE, min.pct = 0.25, logfc.threshold = 0.25)
Cell_crush_markers20 <- Cell_crush_markers %>% group_by(cluster) %>% slice_max(avg_log2FC, n = 20) #make a table of the top 20 expressed genes in each cluster

Cell_crush_markers20 #prints out the table

write.csv(Cell_crush_markers20, file="Cell_crush_markers20.csv") #saves the list of genes as a csv file at your current directory.
```


Based on the marker genes I determine which clusters contain which cell types. In the next bit of code I rename the clusters based on their identity.


```
new_cluster_ids <- c("SGC", "Macrophage", "Fibroblast", "Fibroblast", "Pericyte", "Endothelial","Lymphoid", "Neuron", "Schwann cell", "Myeloid", "Erythrocyte") #The name of the cells need to be written in the order that fits with the clustering. Eg SGC are cluster 0 and Macrophage are cluster 1. You should check that the order is correct.
names(new_cluster_ids) <- levels(Cell_crush)
Cell_crush <- RenameIdents(Cell_crush, new_cluster_ids)
Cell_crush$celltype <- Idents(Cell_crush)
```


To get the other half of figure 1A:


```
DimPlot(Cell_crush, reduction = "umap", label = TRUE, repel = TRUE) + NoLegend()
```


FIGURE 1B Cell\_SNI contains two assays. One called “integrated” with the integrated data used to make the UMAPs and clustering and one called “RNA” which is the unintegrated data.


```
DefaultAssay(Cell_SNI) <- "RNA" #Here the assay is switched to the unintegrated RNA assay
FeaturePlot(Cell_SNI, features = c("Ncmap", "Fabp7", "Cldn5", "Dcn", "Lyz2", "Tubb3", "Rgs5"))
```


FIGURE 1C: To do the comparison between the two datasets shown in figure 1C we first need to convert the Cell\_crush and Cell\_SNI into single cell experiment objects. For more details on this analysis please see https://bioconductor.org/packages/release/bioc/html/scmap.html


```
DefaultAssay(Cell_crush) <- "RNA"
DefaultAssay(Cell_SNI) <- "RNA"

Cell_crush_sce <- as.SingleCellExperiment(Cell_crush)
Cell_SNI_sce <- as.SingleCellExperiment(Cell_SNI)
```


First, we need to prepare the single cell experiment objects:


```
#use gene names as feature symbols
rowData(Cell_SNI_sce)$feature_symbol <- rownames(Cell_SNI_sce)
# remove features with duplicated names
Cell_SNI_sce <- Cell_SNI_sce[!duplicated(rownames(Cell_SNI_sce)), ]
Cell_SNI_sce
```


```
#use gene names as feature symbols
rowData(Cell_crush_sce)$feature_symbol <- rownames(Cell_crush_sce)
# remove features with duplicated names
Cell_crush_sce <- Cell_crush_sce[!duplicated(rownames(Cell_crush_sce)), ]
Cell_crush_sce
```


Next feature selection


```
Cell_crush_sce <- selectFeatures(Cell_crush_sce, suppress_plot = FALSE)
```


```
Cell_crush_sce <- indexCluster(Cell_crush_sce, cluster_col = "ident")
```


We use the scmap cell function to do the projection:


```
set.seed(1)
```


Index


```
Cell_crush_sce <- indexCell(Cell_crush_sce)
```


Projection


```
scmapCell_results <- scmapCell(
  Cell_SNI_sce, 
  list(
    yan = metadata(Cell_crush_sce)$scmap_cell_index
  )
)
```


```
scmapCell_clusters <- scmapCell2Cluster(
  scmapCell_results, 
  list(
    as.character(colData(Cell_crush_sce)$ident)
  )
)
```


```
plot(
  getSankey(
    colData(Cell_SNI_sce)$ident, 
    scmapCell_clusters$scmap_cluster_labs[,"yan"],
    plot_height = 300,
    plot_width = 200,
    colors = c("#00BFC4", "#D89000", "#A3A500", "#39B600", "#00BF7D", "#F8766D", "#00B0F6", "#9590FF", "#E76BF3", "#FF62BC")
  )
)
```


The Sankey plot should pop us a new browser window.

FIGURE 1D: To make the bar plots illustrating the percentage of various cell types shown in figure 1D we first need to calculate the percentages. First we make the bar plot for the Cell\_SNI dataset.


```
prop.table(table(Cell_SNI$orig.ident, Cell_SNI$celltype), margin=1)*100
```


The percentages is placed in the function as values and we create a data frame with the following coloumns: celltype, time and value.


```
df_SNI_per <- data.frame(celltype = rep(c("Schwann cell",  "SGC", "Endothelial", "Macrophage", "Neuron", "Pericyte", "Fibroblast", "Lymphoid", "Myeloid", "Erythrocyte"), each = 3),
                     time = rep(c("Naive", "7 days", "14 days"),10),
                    value = c(42.7, 39.5, 39.7, 33, 25.5, 28.1, 9.1, 7.8, 8.3, 3.3, 9.5, 8.6, 4.1, 8.6, 3.6, 4.1, 3.9, 4.7, 2.7, 2.8, 4.9, 0.3, 1, 1, 0.5, 0.7, 0.6, 0.3, 0.7, 0.6)
                    )

df_SNI_per
```


We take the data frame and turn it into a bar plot.


```
times <- c("Naive", "7 days", "14 days")
ggplot(data=df_SNI_per, aes(x=time, y=value, fill=celltype)) +
  geom_bar(stat="identity")+scale_fill_manual(values=c("#A3A500", "#FF62BC", "#00B0F6", "#9590FF","#39B600", "#E76BF3", "#00BF7D", "#F8766D", "#00BFC4", "#D89000"))+theme_minimal()+scale_x_discrete(limits = times)+theme(legend.text=element_text(size=15), axis.text = element_text(size=15))
```


We do the same for the Cell\_crush dataset


```
prop.table(table(Cell_crush$orig.ident , Cell_crush$celltype), margin=1)*100/2
```


```
df_crush_per <- data.frame(celltype = rep(c("Schwann cell",  "SGC", "Endothelial", "Macrophage", "Neuron", "Pericyte", "Fibroblast", "Lymphoid", "Myeloid", "Erythrocyte"), each = 2),
                     time = rep(c("Naive", "3 days"),10),
                    value = c(5, 3.2, 25.4, 33.3, 7.2, 6.6, 18.2, 23.2, 6.1, 4, 9.4, 6.1, 23.3, 19.7, 2.2, 0.3, 2.1, 1.8, 0.5, 1.7)
                    )

df_crush_per
```


```
times_crush <- c("Naive", "3 days")
ggplot(data=df_crush_per, aes(x=time, y=value, fill=celltype)) +
  geom_bar(stat="identity")+scale_fill_manual(values=c("#A3A500", "#FF62BC", "#00B0F6", "#9590FF","#39B600", "#E76BF3", "#00BF7D", "#F8766D", "#00BFC4", "#D89000"))+theme_minimal()+scale_x_discrete(limits = times_crush)+theme(legend.text=element_text(size=15), axis.text = element_text(size=15))
```


FIGURE 2A-C

Figure 2A-C is a comparison of the SGCs from the different conditions. First we subset the SGCs from the two dataset and filter out possible doublets based on gene detection (nFeature counts).


```
Cell_SNI_SGC <- subset(x=Cell_SNI, idents = "SGC", nFeature_RNA < 3500)
Cell_crush_SGC <- subset(x=Cell_crush, idents = "SGC", nFeature_RNA < 2500)
```


The data subsets are set to the “RNA” assay and a metadata column is added:


```
DefaultAssay(Cell_SNI_SGC) <- "RNA"
DefaultAssay(Cell_crush_SGC) <- "RNA"
Cell_SNI_SGC$orig.data <- "Cell_SNI"
Cell_crush_SGC$orig.data <- "Cell_crush"
```


The datasets are merged:


```
Cell_SGC <- merge(Cell_crush_SGC, y=Cell_SNI_SGC)
```


Next the data subsets need to be integrated:


```
# split the dataset into a list of two seurat objects (stim and CTRL)
ifnb.list <- SplitObject(Cell_SGC, split.by = "orig.ident")

# normalize and identify variable features for each dataset independently
ifnb.list <- lapply(X = ifnb.list, FUN = function(x) {
    x <- NormalizeData(x)
    x <- FindVariableFeatures(x, selection.method = "vst", nfeatures = 2000)
})

# select features that are repeatedly variable across datasets for integration
features <- SelectIntegrationFeatures(object.list = ifnb.list)
```


Perform integration:


```
SNI_anchors <- FindIntegrationAnchors(object.list = ifnb.list, anchor.features = features, dim=1:20, k.filter=100)
# this command creates an 'integrated' data assay
Cell_SGC_integrated <- IntegrateData(anchorset = SNI_anchors, dim=1:20)
```


We perform the visualization and clustering:


```
# specify that we will perform downstream analysis on the corrected data note that the original
# unmodified data still resides in the 'RNA' assay
DefaultAssay(Cell_SGC_integrated) <- "integrated"

# Run the standard workflow for visualization and clustering
Cell_SGC_integrated <- ScaleData(Cell_SGC_integrated, verbose = FALSE)
Cell_SGC_integrated <- RunPCA(Cell_SGC_integrated, npcs = 30, verbose = FALSE)
Cell_SGC_integrated <- RunUMAP(Cell_SGC_integrated, reduction = "pca", dims = 1:20)
Cell_SGC_integrated <- FindNeighbors(Cell_SGC_integrated, reduction = "pca", dims = 1:20)
Cell_SGC_integrated <- FindClusters(Cell_SGC_integrated, resolution = 0.2)
```


Next we need a new metadata column where the data set is named based on being injured or uninjured.


```
#The idents are set to be orig.ident which is inj1, inj2, con1, con2, 7 days, 14 days and naive)
Cell_SGC_integrated <- SetIdent(Cell_SGC_integrated, value=Cell_SGC_integrated@meta.data$orig.ident)
#We make a vector with the new corresponding names
condition <- c("Injury", "Injury", "Naive", "Naive", "Injury", "Injury", "Naive")
names(condition) <- levels(Cell_SGC_integrated)
Cell_SGC_integrated <- RenameIdents(Cell_SGC_integrated, condition)
Cell_SGC_integrated$condition <- Idents(Cell_SGC_integrated)
```


```
# Visualization
p1 <- DimPlot(Cell_SGC_integrated, reduction = "umap", group.by = "seurat_clusters", label=TRUE, label.size = 7, pt.size = 1)
p2 <- DimPlot(Cell_SGC_integrated, reduction = "umap", group.by = "condition", pt.size = 1)
p3 <- DimPlot(Cell_SGC_integrated, reduction = "umap", group.by = "orig.data", pt.size = 1)
p1 + p2 + p3
```


FIGURE 3 Next we move on to figure 3 which look at the Cell\_Culture data. The dataset can be downloaded from GEO on accession GSE188971


```
Cell_culture <- readRDS("C:/Users/your/path/Cell_culture_data.rds") #Adjust the path to fit your system
```


Figure 3A:


```
DimPlot(Cell_culture, reduction = "umap", label = TRUE, pt.size = 0.5, group.by = "celltype", label.size = 5)
```


Figure 3B:


```
DefaultAssay(Cell_culture) <-  "RNA"
FeaturePlot(Cell_culture, features=c("Fabp7", "Kcnj10", "Bcas1", "Prx", "Ncmap", "Tubb3", "Dcn", "Lyz2"))
```


FIGURE 4 In figure 4 we move on to the joint analysis between the Cell\_culture and the Cell\_SNI datasets.The joint analysis is done by merging and integrating the Cell\_SNI and Cell\_culture datasets. The seurat object can be downloaded here: GSE188971


```
Cell_SNI_culture <- readRDS("C:/Users/your/path/Cell_SNI_culture_data.rds") #Adjust the path to fit your system
```


Figure 4A+B


```
# Visualization
p4 <- DimPlot(Cell_SNI_culture, reduction = "umap",label = TRUE, repel = TRUE)
p5 <- DimPlot(Cell_SNI_culture, reduction = "umap", group.by = "orig.ident")

p4 + p5
```


Figure 4C In Figure 4C we show how the annotation from the joint analysis look when only displaying the Cell\_culture cells in an UMAP:

I subset the data to only have the cells from the Cell\_culture dataset:


```
Cell_culture_sub <- subset(Cell_SNI_culture, subset = dataset == "Cell_culture")
DefaultAssay(Cell_culture_sub) <- "RNA"
```


Next I normalize and scale the data:


```
Cell_culture_sub <- NormalizeData(Cell_culture_sub)
Cell_culture_sub <- ScaleData(Cell_culture_sub)
```


Next I integrate them:


```
# split the dataset into a list of two seurat objects (stim and CTRL)
ifnb.list <- SplitObject(Cell_culture_sub, split.by = "orig.ident")

# normalize and identify variable features for each dataset independently
ifnb.list <- lapply(X = ifnb.list, FUN = function(x) {
    x <- NormalizeData(x)
    x <- FindVariableFeatures(x, selection.method = "vst", nfeatures = 2000)
})

# select features that are repeatedly variable across datasets for integration
features <- SelectIntegrationFeatures(object.list = ifnb.list)
```


Perform integration:


```
Cell_culture_anchors <- FindIntegrationAnchors(object.list = ifnb.list, anchor.features = features, dim=1:20)
# this command creates an 'integrated' data assay
Cell_culture_sub <- IntegrateData(anchorset = Cell_culture_anchors, dim=1:20)
```


```
# specify that we will perform downstream analysis on the corrected data note that the original
# unmodified data still resides in the 'RNA' assay
DefaultAssay(Cell_culture_sub) <- "integrated"

# Run the standard workflow for visualization and clustering
Cell_culture_sub <- ScaleData(Cell_culture_sub, verbose = FALSE)
Cell_culture_sub <- RunPCA(Cell_culture_sub, npcs = 30, verbose = FALSE)
Cell_culture_sub <- RunUMAP(Cell_culture_sub, reduction = "pca", dims = 1:20)
Cell_culture_sub <- FindNeighbors(Cell_culture_sub, reduction = "pca", dims = 1:20)
Cell_culture_sub <- FindClusters(Cell_culture_sub, resolution = 0.18)
```


```
p6 <- DimPlot(Cell_culture_sub, reduction = "umap", group.by = "celltype_int", label = TRUE, repel = TRUE, label.size = 5) 
p6
```


The UMAP generated here might look a bit different to the once in the paper. This is because UMAPs have a random component. The clusters are reproducible, so it is only the UMAP visualization that varies.

Figure 4D


```
p7 <- DimPlot(Cell_SNI_culture, reduction = "umap", split.by = "dataset", label = TRUE, label.size = 5) + NoLegend()
p7
```


Figure 4E


```
DefaultAssay(Cell_SNI_culture) <- "RNA"
FeaturePlot(Cell_SNI_culture, features=c("Fabp7", "Kcnj10", "Top2a", "Mki67", "Ncmap", "Bcas1", "Prx"))
```


Figure 4F: In this figure we compare the gene expression in the joint analysis with gene expression in the developing nervous system from Furlan et al., Science, 2017.

First we need to download the data from Furlan et al. Download the GSE99933\_E12.5\_counts.txt.gz file from Gene Expression Omnibus (GEO) database at accession number GSE99933 under supplementary files. Extract the data so your data file is in the .txt format.


```
precursor_data <- read.delim("C:/Users/your/path/E12.5_counts.txt", header=TRUE) #Adjust the path to fit your system
#Change the data to a Seurat object
precursor <- CreateSeuratObject(counts=precursor_data)
precursor
```


There is no meta data connected to the downloaded counts so we have to do the analysis:


```
precursor <- NormalizeData(precursor)
precursor <- FindVariableFeatures(precursor, selection.method = "vst", nfeatures = 2000)
all.genes <- rownames(precursor)
precursor <- ScaleData(precursor, features = all.genes)
precursor <- RunPCA(precursor, features = VariableFeatures(object = precursor))
precursor <- FindNeighbors(precursor, dims = 1:14)
precursor <- FindClusters(precursor, resolution = 0.25)
precursor <- RunUMAP(precursor, dims = 1:14)
DimPlot(precursor, reduction = "umap")
```


To find out which cluster is which cell type we check the markers from the Furlan et al paper:


```
FeaturePlot(precursor, feature=c("Sox10", "Th", "Foxq1", "Chgb"))
```


Based on this we can rename the clusters:


```
new.cluster.ids <- c("Bridge cells", "SCPs", "Chromaffin cells", "Sympathoblasts")
names(new.cluster.ids) <- levels(precursor)
precursor <- RenameIdents(precursor, new.cluster.ids)
DimPlot(precursor, reduction = "umap", label = TRUE, pt.size = 2)
```


Next we turn the object into a Single Cell experiment object:


```
precursor_exp <- as.SingleCellExperiment(precursor)
```


I also change the Cell\_SNI\_culture dataset into a Single Cell experiment object:


```
Cell_SNI_culture_exp <- as.SingleCellExperiment(Cell_SNI_culture, assay="RNA")
```


We are now ready to run the comparison analysis with SingleR:


```
matched_precursor <- matchReferences(precursor_exp, Cell_SNI_culture_exp, precursor_exp$ident, Cell_SNI_culture$celltype_int) #This command takes 5 min or so to run. 
pheatmap::pheatmap(matched_precursor, col=viridis::plasma(100))
```


FIGURE 5 Next we move on to figure 5 were we do a closer comparison of the SGCs from the Cell\_culture and the Cell\_SNI dataset.


```
Cell_SNI_culture_SGC <- subset(Cell_SNI_culture, subset = celltype_int =="SGC")
Cell_SNI_culture_SGC
```


Integrate:


```
# split the dataset into a list of two seurat objects (stim and CTRL)
ifnb.list <- SplitObject(Cell_SNI_culture_SGC, split.by = "orig.ident")

# normalize and identify variable features for each dataset independently
ifnb.list <- lapply(X = ifnb.list, FUN = function(x) {
    x <- NormalizeData(x)
    x <- FindVariableFeatures(x, selection.method = "vst", nfeatures = 2000)
})

# select features that are repeatedly variable across datasets for integration
features <- SelectIntegrationFeatures(object.list = ifnb.list)
```


Perform integration:


```
Cell_culture_anchors <- FindIntegrationAnchors(object.list = ifnb.list, anchor.features = features, dim=1:20)
# this command creates an 'integrated' data assay
Cell_SNI_culture_SGC <- IntegrateData(anchorset = Cell_culture_anchors, dim=1:20)
```


Visualize:


```
# specify that we will perform downstream analysis on the corrected data note that the original
# unmodified data still resides in the 'RNA' assay
DefaultAssay(Cell_SNI_culture_SGC) <- "integrated"

# Run the standard workflow for visualization and clustering
Cell_SNI_culture_SGC <- ScaleData(Cell_SNI_culture_SGC, verbose = FALSE)
Cell_SNI_culture_SGC <- RunPCA(Cell_SNI_culture_SGC, npcs = 30, verbose = FALSE)
Cell_SNI_culture_SGC <- RunUMAP(Cell_SNI_culture_SGC, reduction = "pca", dims = 1:20)
Cell_SNI_culture_SGC <- FindNeighbors(Cell_SNI_culture_SGC, reduction = "pca", dims = 1:20)
Cell_SNI_culture_SGC <- FindClusters(Cell_SNI_culture_SGC, resolution = 0.2)
```


Figure 5A


```
# Visualization
my_col2 <-c("14 days"="#00B0F6", "7 days"="#00B0F6", "Naive"="#00B0F6", "A"="#F8766D", "B"="#F8766D")
p8 <- DimPlot(Cell_SNI_culture_SGC, reduction = "umap", group.by = "orig.ident", cols = my_col2)
p9 <- DimPlot(Cell_SNI_culture_SGC, reduction = "umap",label = TRUE, repel = TRUE, label.size = 7)
p8 + p9
```


The UMAP generated here might look a bit different to the once in the paper. This is because UMAPs have a random component. The clusters are reproducible, so it is only the UMAP visualization that varies.

Figure 5B

In this figure we compare the gene expression in the clusters identified in figure 5A with cell types in the developing nervous system. First we need to turn the Cell\_SNI\_culture\_SGC into a Single Cell experiment:


```
SGC_exp <- as.SingleCellExperiment(Cell_SNI_culture_SGC, assay="RNA")
```


We can now perform the comparison analysis:


```
matched_precursor <- matchReferences(precursor_exp, SGC_exp, precursor_exp$ident, SGC_exp$ident)#This code takes about 5 min to run
pheatmap::pheatmap(matched_precursor, col=viridis::plasma(100))
```
